# ECM1 Attenuates Hepatic Fibrosis by Interfering with Mediators of Latent TGF-β1 Activation

**DOI:** 10.1101/2023.12.12.571289

**Authors:** Frederik Link, Yujia Li, Jieling Zhao, Stefan Munker, Weiguo Fan, Zeribe C. Nwosu, Ye Yao, Shanshan Wang, Chenjun Huang, Roman Liebe, Seddik Hammad, Hui Liu, Chen Shao, Chunfang Gao, Bing Sun, Natalie J. Török, Huiguo Ding, Matthias P. A. Ebert, Honglei Weng, Peter ten Dijke, Dirk Drasdo, Steven Dooley, Sai Wang

## Abstract

**Objective:** Extracellular Matrix Protein 1 (ECM1) serves as a gatekeeper of hepatic fibrosis by maintaining transforming growth factor-β1 (TGF-β1) in its latent form. ECM1 knockout (KO) causes latent (L) TGF-β1 activation, resulting in hepatic fibrosis with rapid mortality. In chronic liver disease (CLD), ECM1 decreases with increasing CLD severity. We investigate the regulatory role of ECM1 in TGF-β1 bioavailability and its impact on CLD progression.

**Design:** RNAseq was performed to analyze hepatic gene expression. Functional assays were performed using hepatic stellate cells (HSCs), *Ecm1*-KO and *Fxr*-KO mice, patient liver tissue, and computer simulations.

**Results:** Expression of LTGF-β1 activators, including thrombospondins (TSPs), ADAMTS proteases, and matrix metalloproteinases (MMPs) increased along with pro-fibrotic gene expression in liver tissue of *Ecm1*-KO mice. In HSCs, overexpression of ECM1 prevented TSP-1-, ADAMTS1-, and MMP-2/9-mediated LTGF-β1 activation. *In vitro* interaction assays demonstrated that ECM1 inhibited LTGF-β1 activation by interacting with TSP-1 and ADAMTS1 via their respective, intrinsic KRFK or KTFR amino acid sequences, and by suppressing MMP-2/9 proteolytic activity. In mice, ECM1 overexpression attenuated KRFK-induced LTGF-β1 activation, while KTFR treatment reversed *Ecm1*-KO- and *Fxr*-KO-mediated liver injury. In patients with CLD, ECM1 expression was inversely correlated with TSP-1, ADAMTS1, MMP-2/9 expression and LTGF-β1 activation. And these results were complemented by a computational compartment model representing the key network of cellular phenotypes and predicted interactions in liver fibrogenesis.

**Conclusion:** Our findings underscore the hepatoprotective effect of ECM1, which interferes with mediators of LTGF-β1 activation, suggesting ECM1 or its representative peptide as potential anti-fibrotic therapies in CLD.

**What is already known on this topic?:**

➢ ECM1 expression is negatively correlated with CLD progression.
➢ ECM1 maintains liver homeostasis by keeping TGF-β1 latency.

**What this study adds?:**

➢ ECM1 inhibits LTGF-β1 activation through interfering with key activators, including TSP-1, ADAMTS1, MMP-2, and MMP-9.
➢ ECM1 interacts with TSP-1 and ADAMTS1 via their respective, intrinsic KRFK or KTFR amino acid motifs, and suppresses MMP-2/9 proteolytic activity.
➢ *In vivo*, ECM1 overexpression mitigates KRFK peptide-induced LTGF-β1 activation, while KTFR peptide rescues *Ecm1*-KO- and *Fxr*-KO-induced liver injury.
➢ ECM1 expression inversely correlates with TSP-1, ADAMTS1, MMP-2/9 expression and LTGF-β1 activation in CLD patients.

**How might this study affect research, practice or policy?:**

➢ Considering severe adverse effects associated with anti-fibrotic treatments utilizing TGF-β1 receptor inhibitors, our findings indicate that restoration of ECM1 expression or phenocopying peptides might represent a novel and safe route to urgently needed anti-fibrotic therapies in CLD.

## Introduction

Hepatic fibrosis is the main clinical outcome of CLD, paving the ground for the development of liver cirrhosis and hepatocellular carcinoma (HCC) [1]. Fibrotic diseases constitute an increasing global health burden, with no specific treatments available [2]. Extracellular matrix protein 1 (ECM1), a secreted glycoprotein [3], is essential for the maintenance of tissue homeostasis [4, 5]. In the liver, a gradual decrease of ECM1 expression has been associated with progressive hepatic fibrosis [6]. Complete depletion of ECM1 in mice results in severe and lethal liver fibrosis accompanied by drastically upregulated transforming growth factor-β (TGF-β)1 signaling [6]. ECM1 was also discovered to be the third most abundant gene encoding a secreted protein in quiescent hepatic stellate cells (HSC) and shown to be downregulated during their transdifferentiation to fibrogenic myofibroblasts [7], suggesting that quiescent HSCs are important providers of ECM1 to maintain hepatic homeostasis.

Targeting TGF-β1 signaling is considered a risky therapeutic strategy in many organs due to its heterogenous and contextual cellular functions [8, 9]. This includes the liver, where loss or complete inhibition of TGF-β1 results in developmental disorders, deleterious multi-organ inflammation, and early postnatal death [10], while excessive TGF-β1 signaling in CLD causes fibrosis progression and tumor metastasis [11, 12, 13]. Due to its broad range of functions and effects, the bioavailability of TGF-β1 is tightly controlled [14]. Upon intracellular synthesis, TGF-β1 is secreted and stored covalently attached to ECM molecules as an inactive protein complex termed latent TGF-β1 (LTGF-β1) [15]. Activation of LTGF-β1 involves a range of mechanisms [14], including integrins [16, 17], proteases [18], reactive oxygen species (ROS) [19], mechanical force [16, 20], and various other physical and chemical cues [21, 22]. Progressive hepatic fibrosis is driven by increasing levels of active TGF-β1 [23, 24]. Targeting the active TGF-β1 molecule through direct inhibition has not proven feasible due to severe adverse effects. Therefore, our current research aimed at uncovering the regulatory mechanisms behind the bioavailability of TGF-β1 to develop safer anti-fibrotic strategies [14]. Specifically, we examined whether and how ECM1 prevents LTGF-β1 activation *in vitro* and *in vivo*. Moreover, we complemented our analysis by a computational model that implements major cell types and inter-/intracellular communications that may be affected by changes in the concentration of ECM1.

## Results

### Ecm1 depletion induces mediators able to activate LTGF-β1 and hepatic fibrogenesis

*Ecm1* depletion (*Ecm1*-KO) in mice leads to hepatic fibrosis and rapid mortality at the age of 8-12 weeks [6]. To investigate mechanisms involved in *Ecm1*-KO-induced liver injuries, liver tissue from WT and *Ecm1*-KO mice at 2, 5, and 8 weeks of age (n = 3) was collected for RNAseq analyses. As TGF-β1 is already known to be the central mediator of liver injury in *Ecm1*-KO mice [6], we analyzed changes in gene transcription specifically in the context of this cytokine. Consequently, we sought for the expression of candidate factors that are capable of LTGF-β1 activation in young (2-weeks-old) *Ecm1*-KO mice. We discovered that *Ecm1*-KO mice expressed considerably higher levels of thrombospondins (*Tsps*) and several *Adamts* proteases, including *Adamts4*, *Adamtsl2*, *Thbs1* (*Tsp1*), *Adamts9*, *Thbs2* (*Tsp2*), *Adamts2*, *Adamts1* **(Figure 1A)**, some of which are known to activate LTGF-β1 or interact with the latency associated peptide (LAP) of LTGF-β1 [25, 26]. Additionally, numerous matrix metalloproteases (*Mmps)* known to activate LTGF-β1 *in vitro* [27] including *Mmp-2*, *Mmp-8*, *Mmp-13*, *Mmp-9* were upregulated in *Ecm1*-KO mice **(Figure 1A)**. The chord diagram demonstrated that these elevated molecues are components of the extracellular matrix (ECM) and implicated in ECM (re-)organization due to their peptidase or proteolytic activity **(Figure 1B)**, hence supporting their critical role in tissue remodeling during fibrosis. Furthermore, networking analyses showed that the majority of these Tsps, Adamts proteases, and Mmps share protein domains, while some of them also show co-localization, physical interaction, co-regulated pathways, and genetic interactions **(Figure 1C)**. Knowing its direct effect on integrin activity, we hypothesized that ECM1 may be a modulator of these LTGF-β1 activators, eventually possibly, through binding to shared protein domains. To further investigate the correlation of these factors with LTGF-β1 activation in the setting of *Ecm1* depletion, we analyzed their participation in functional pathways and found notable upregulation in endodermal cell differentiation, cell migration, and vascular smooth muscle cell proliferation in *Ecm1*-KO mice, all well-known as regulated by means of TGF-β signaling **(Figure 1D, E)**, suggesting that all upregulated factors are substantially involved in LTGF-β1 activation when ECM1 is depleted. Indeed, when *Ecm1* was knocked out, expression of Tsp-1, Adamts1, Mmp-2 and -9 were all increased on mRNA and protein level **(Figure 1F, G)**.

**Figure 1.**
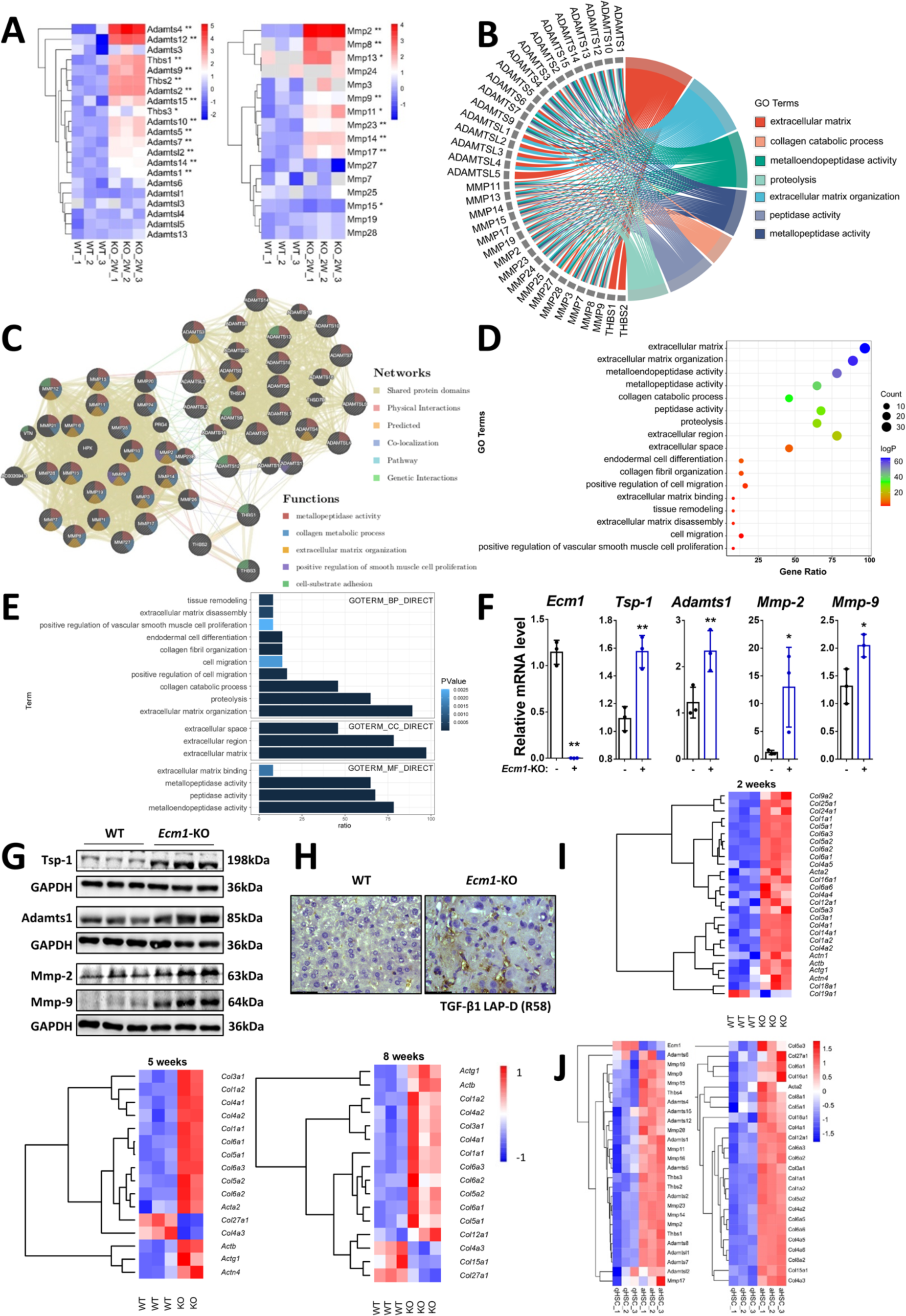
ECM1 depletion causes the upregulation of mediators of LTGF-β activation and severe fibrogenesis in liver. **(A)** RNAseq analysis of the expression of *Mmps*, *Tsps*, and *Adamtss* in 2 weeks old *Ecm1*-KO as compared to *WT* mice. **(B)** Chord diagram, illustrating the main biological processes the upregulated proteases are involved in. **(C)** Interconnectivity of the upregulated MMps, Tsps, and Adamtss. **(D, E)** GO pathways influenced by *Ecm1*-KO-induced proteases. **(F, G)** Relative mRNA and protein levels of Tsp-1, Adamts1, Mmp-2, and Mmp-9 in liver tissue from WT and *Ecm1*-KO mice. **(H)** Representative images of IHC stainings for TGF-β1 LAP-D (R58) in liver tissue from *Ecm1*-KO and *WT* mice. Scale bar, 43.5 µm. **(I)** RNA-Seq analysis, demonstrating the upregulation of fibrotic genes in *Ecm1*-KO mice at the age of 2, 5 and 8 weeks. **(J)** mRNA expression of *Ecm1*, LTGF-β1 activators, and fibrotic markers in quiescent and activated HSCs isolated from control or CCl_4_-treated mice (n=3), extracted from GEO DataSets GSE149508. For RT-qPCR, human *PPIA* was used as an endogenous control. For Western blotting, GAPDH was used as a loading control. P-values were calculated by unpaired Student’s t-test. Bars represent mean ± SD. *p<0.05; **p<0.01.

Next, we employed an antibody that selectively recognizes the TGF-β1 latency associated peptide (LAP) breakdown product (LAP-D R58), which is retained in the ECM following LTGF-β1 activation. Immunohistochemistry (IHC) staining showed that, compared to WT mice, there was enhanced LTGF-β1 activation in the liver tissue of *Ecm1*-KO mice, with LAP-D R58 expression detected in the (peri-)sinusoidal area **(Figure 1H)**. Consistent with the profibrogenic effects of TGF-β1, many genes encoding for prominent matrisomal proteins, such as collagen type 1 (*Col1α1*, *Col1α2*), collagen type 3 (*Col3α1*), collagen type 5 (*Col5α1*, *Col5α2*, *Col5α3*), collagen type 6 (*Col6α1*, *Col6α2*, *Col6α3*), and the HSC activation marker *Acta2* were significantly upregulated in *Ecm1*-KO mice, and remained elevated over the investigated time course **(Figure 1I)**. Intriguingly, *Col27a1*, *Col4a3*, and *Col15a* distinguish themselves from the aforementioned fibrillous collagens, as they are important structural components of the ECM in healthy livers and were decreased in *Ecm1*-KO mice at 5 or 8 weeks, further indicating ECM1’s substantial role in liver tissue homeostasis **(Figure 1I, bottom left, 2 panels)**. Given that HSCs were identified as the primary producers of TGF-β1 and its activators, we analyzed mRNA expression for our target genes in primary quiescent and activated HSCs isolated from control and CCl_4_-treated mice (n=3) from the Gene Expression Omnibus (GEO) DataSets GSE149508 [28]. Specifically, compared to quiescent HSCs, *Ecm1* exhibited a noteworthy downregulation, whereas *Tsps*, *Adamts proteases*, *Mmps*, and fibrotic *collagens* demonstrated substantial upregulation in activated HSCs **(Figure 1J)**, in line with the results in *Ecm1*-KO mice.

To summarise, ECM1 depletion causes notable upregulation of known activators of LTGF-β1, including TSPs, ADAMTS proteases, and MMPs, which may subsequently mediate activation of large amounts of LTGF-β1 deposited in the ECM, causing the lethal hepatic fibrosis seen in *Ecm1*-KO mice.

### ECM1 inhibits αvβ6 integrin-mediated LTGF-β1 activation

Our previous study showed that ECM1 inhibits αvβ6 integrin-mediated LTGF-β1 activation by interference with its binding to LTGF-β1’s RGD motif [6]. To mechanistically study the inhibitory effect of ECM1 on LTGF-β1 activation in further detail, we set up an *in vitro* system comprising HSCs and the TGF-β/SMAD signaling reporter cell line MFB-F11, in which the active TGF-β in the supernatant of HSCs can be measured by incubating the conditioned media with MFB-F11 cells and performing a SEAP (secreted alcaline phosphatase) activity assay **(Figure 2A)**, which is more sensitive than a TGF-β1 ELISA (**Figure S1A, B**). For functional proof of the assay, we tested the cells with αvβ6 integrin as a positive control for LTGF-β1 activation. As anticipated, pre-treatment of LX-2 HSCs and primary human (ph) HSCs that predominantly produce TGF-β1 [29, 30] with 100ng/ml recombinant ECM1 (rECM1) or transfection with an ECM1 plasmid (V-ECM1; LX-2 HSCs) (pcDNA3.1 was transfected as a control plasmid), before adding αvβ6 integrin, both significantly reduced TGF-β signaling reporter activation in MFB-F11 cells, as compared to controls **(Figure 2B)**. 100 ng/ml rECM1 was selected from an experiment using different concentrations of rECM1 and LTGF-β1 activation by SEAP assay (**Figure S1C**). ECM1’s successful overexpression was checked by RT-qPCR, immunoblotting, and IF staining (**Figure S2**). Our data suggested that ectopic, ECM1-dependent inhibition of αvβ6 integrin-mediated LTGF-β1 activation can be robustly documented with the SEAP assay. In line, mRNA and protein expression of hepatic fibrosis markers *ACTA2*/α-SMA, COL1A1, COL3A1, and TIMP1 were decreased ECM1-dependently **(Figure 2C, D)**. Lastly, the inhibitory effect of ECM1 on αvβ6 integrin-mediated LTGF-β1 activation was detected by measuring the TGF-β1 LAP-D (R58) fragment using IF staining and quantitative image analysis **(Figure 2E)**.

**Figure 2.**
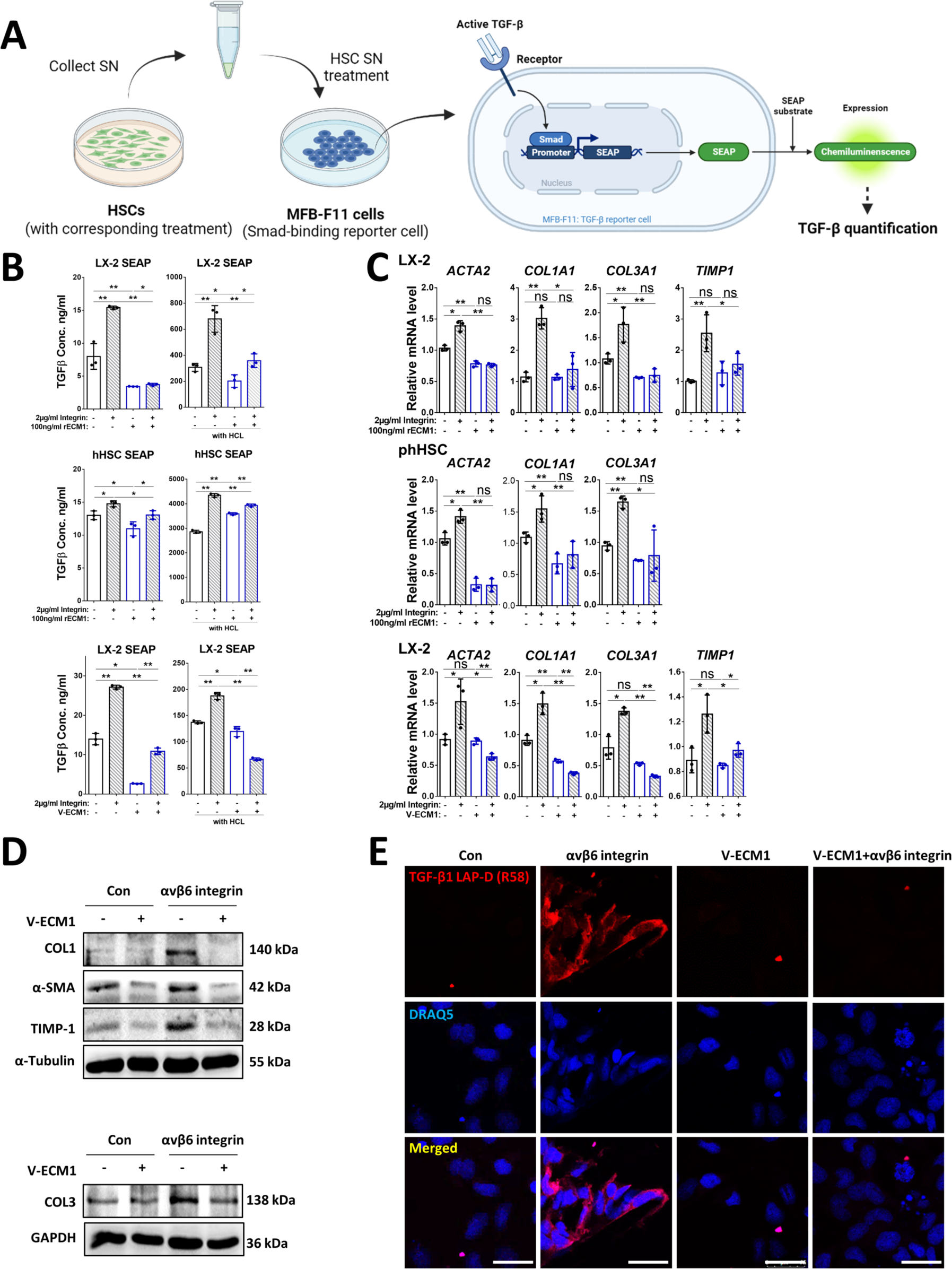
ECM1 inhibits αvβ6 integrin-mediated LTGF-β1 activation. **(A)** Illustration of SEAP activity assay used with the MFB-F11 TGF-β signaling reporter cell line (Created with BioRender.com). **(B)** The concentration of active and total TGF-β (ng/ml) measured by SEAP activity assay from conditioned LX-2 HSCs SN and phHSCs SN incubated w/o rECM1/V-ECM1 and/or αvβ6 integrin. **(C)** Relative mRNA levels of *ACTA2, COL1A1, COL3A1*, and *TIMP1* in LX-2 HSCs and phHSCs incubated w/o rECM1/V-ECM1 and/or αvβ6 integrin. **(D)** Protein levels of COL1, COL3, α-SMA, and TIMP1 in LX-2 HSCs incubated with/without V-ECM1 and/or αvβ6 integrin. **(E)** Representative images of IF stainings for TGF-β1 LAP-D (R58) in LX-2 HSCs incubated with/without V-ECM1 and/or αvβ6 integrin. DRAQ5 was used for nuclear staining. Scale bar, 25 µm. For RT-qPCR, human PPIA was used as an endogenous control. P-values were calculated by unpaired Student’s t-test. Bars represent mean ± SD. *p<0.05; **p<0.01. For Western blotting, GAPDH or α-Tubulin were used as a loading control.

### ECM1 inhibits TSP-1/ADAMTS1/MMP-2/MMP-9-mediated LTGF-β1 activation

Directed by our gene expression data **(Figure 1A, F, G)** and previous literature, we selected four activating mediators (namely, TSP-1, ADAMTS1, MMP-2, and MMP-9) to further study whether ECM1 can prevent LTGF-β1 activation in such a broad and overarching manner. LX-2 HSCs and phHSCs were incubated with rECM1 or transfected with V-ECM1 (LX-2 HSCs) followed by treatment with/without TSP-1, ADAMTS1, MMP-2, or MMP-9 (pcDNA3.1 was used as a control). SEAP activity assay results exemplified that incubation with the activators increased the concentration of active TGF-β1 in conditioned supernatants of LX-2 HSCs as well as phHSCs. LTGF-β1 activation was completely abrogated by prior treatment with rECM1 or upon ECM1 overexpression, assessed for active TGF-β in the supernatant by SEAP assays **(Figure 3A)**. In line with these findings, the expression of fibrosis markers on mRNA level induced by TSP1, ADAMTS1, MMP-2, or MMP-9, including *ACTA2*, *COL1A1*, *COL3A1*, or *TIMP1*, were blunted when rECM1 or V-ECM1 were administered in advance to phHSCs or LX-2 HSCs **(Figure 3B-D)**. We also confirmed ECM1’s inhibitory effect by showing that it reduced expression of fibrosis markers on protein level, compared to the groups treated with one of the four activators, as tested by immunoblotting **(Figure 3E)**. The findings were further underscored by the observed loss of TGF-β1 LAP-D (R58) immunofluorescence, when ECM1 was overexpressed in presence of TSP-1, ADAMTS1, MMP-2, or MMP-9 in LX-2 HSCs **(Figure 3F)**.

**Figure 3.**
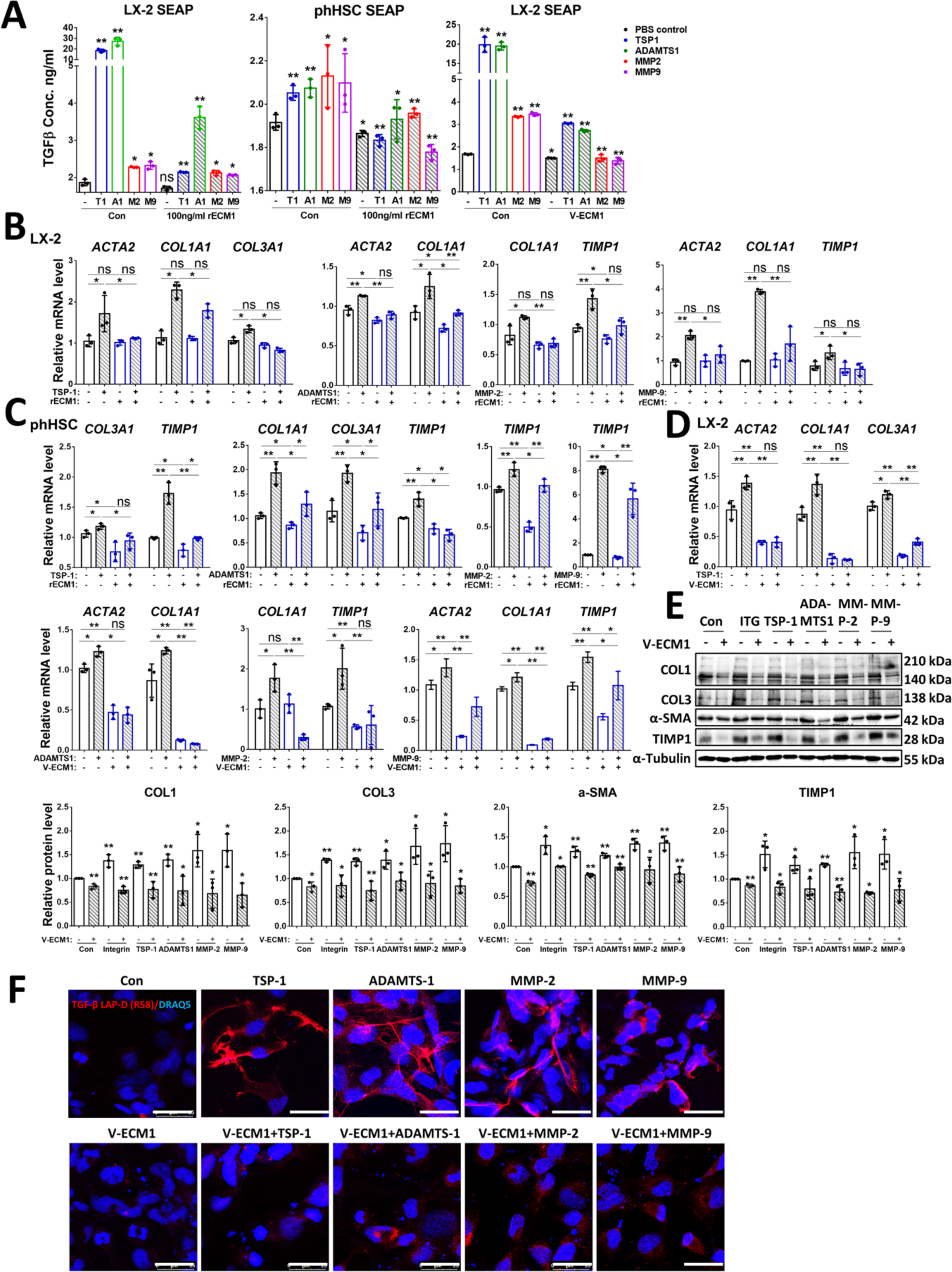
ECM1 inhibits TSP-1/ADAMTS1/MMP-2/MMP-9-mediated LTGF-β1 activation. **(A)** The concentration of active TGF-β (ng/ml) measured by SEAP activity assay from conditioned LX-2 HSCs SN and phHSCs SN incubated w/o TSP-1/ADAMTS1/MMP-2/MMP-9 and/or rECM1/V-ECM1. **(B-D)** Relative mRNA levels of *ACTA2, COL1A1*, *COL3A1*, and *TIMP1* in LX-2 HSCs and phHSCs incubated with/without rECM1/V-ECM1 and/or TSP-1/ADAMTS1/MMP-2/MMP-9. **(E)** Protein levels of COL1, COL3, α-SMA, and TIMP1 in LX-2 HSCs incubated with/without rECM1/V-ECM1 and/or Integrin, TSP-1, ADAMTS1, MMP-2, or MMP-9. **(F)** Representative images of IF stainings for TGF-β1 LAP-D (R58) in LX-2 HSCs incubated with/without V-ECM1 and/or TSP-1/ADAMTS1/MMP-2/MMP-9. DRAQ5 was used for nuclear staining. Scale bar, 25 µm. For RT-qPCR, human PPIA was used as an endogenous control. P-values were calculated by unpaired Student’s t-test. Bars represent mean ± SD. *p<0.05; **p<0.01. For Western blotting, α-Tubulin was used as a loading control. Quantification of protein expression was performed using Image J (National Institute of Health, Bethesda, MD).

### ECM1 inhibits LTGF-β1 activation via interaction with TSP-1/ADAMTS1’s motifs KRFK/KTFR and modulation of MMP-2/9’s proteolytic activity

To confirm the role of ECM1 as a direct and broad inhibitor of multiple LTGF-β1 activators and to exclude the presence of confounding effects from other TGF-β isoforms or other relevant signals in HSCs, we incubated MFB-F11 cells with 4ng/ml recombinant (r)LTGF-β1, followed by treatment with each of the four LTGF-β1 activating mediator, alone or together with rECM1. Again, the presence of rECM1 prevented rLTGF-β1 activation by TSP-1-, ADAMTS1-, MMP-2 and MMP-9, as indicated by the *in vitro* SEAP assay results **(Figure 4A)**. To delve further into how ECM1 inhibits LTGF-β1 activation induced by the above mediators, we performed protein-protein and protein-peptide interaction studies. Immunoprecipitation (IP) analyses illustrated that ECM1 interacted with TSP-1, ADAMTS1, and mature forms of MMP-2 and MMP-9 in LX-2 HSCs that were cultured for 72h **(Figure 4B)**. *In vitro* pull-down assays using purified recombinant proteins confirmed that ECM1 directly bound to TSP-1 (**Figure 4C**), ADAMTS1 (**Figure 4D**), MMP-2 (**Figure 4E**), and MMP-9 (**Figure 4F**). The respective whole immunoblotting membranes can be found in **Figures S3A-D**. **Figure S3E** demonstrated that ECM1 directly interacted with rLTGF-β1 in the *in vitro* pull-down assay. However, it did not affect rLTGF-β1 activation (**Figure 4A, G**, the first and third column in each histogram) thus, ruling out any direct inhibitory effect of ECM1 on LTGF-β1 activation. Incubation of MFB-F11 cells with rECM1 and active TGF-β1 did not affect TGF-β1-induced SEAP signal, which suggests that ECM1 does not directly antagonize active TGF-β1 (**Figure S3F**). Additionally, the Alphafold3 system [31] provided structure predictions on the potential localization of interactions between ECM1 and TSP-1, ADAMTS1, MMP-2, MMP-9, or LTGF-β1 (**Figure S4A-E**), using their full-length target proteins as input sequences.

**Figure 4.**
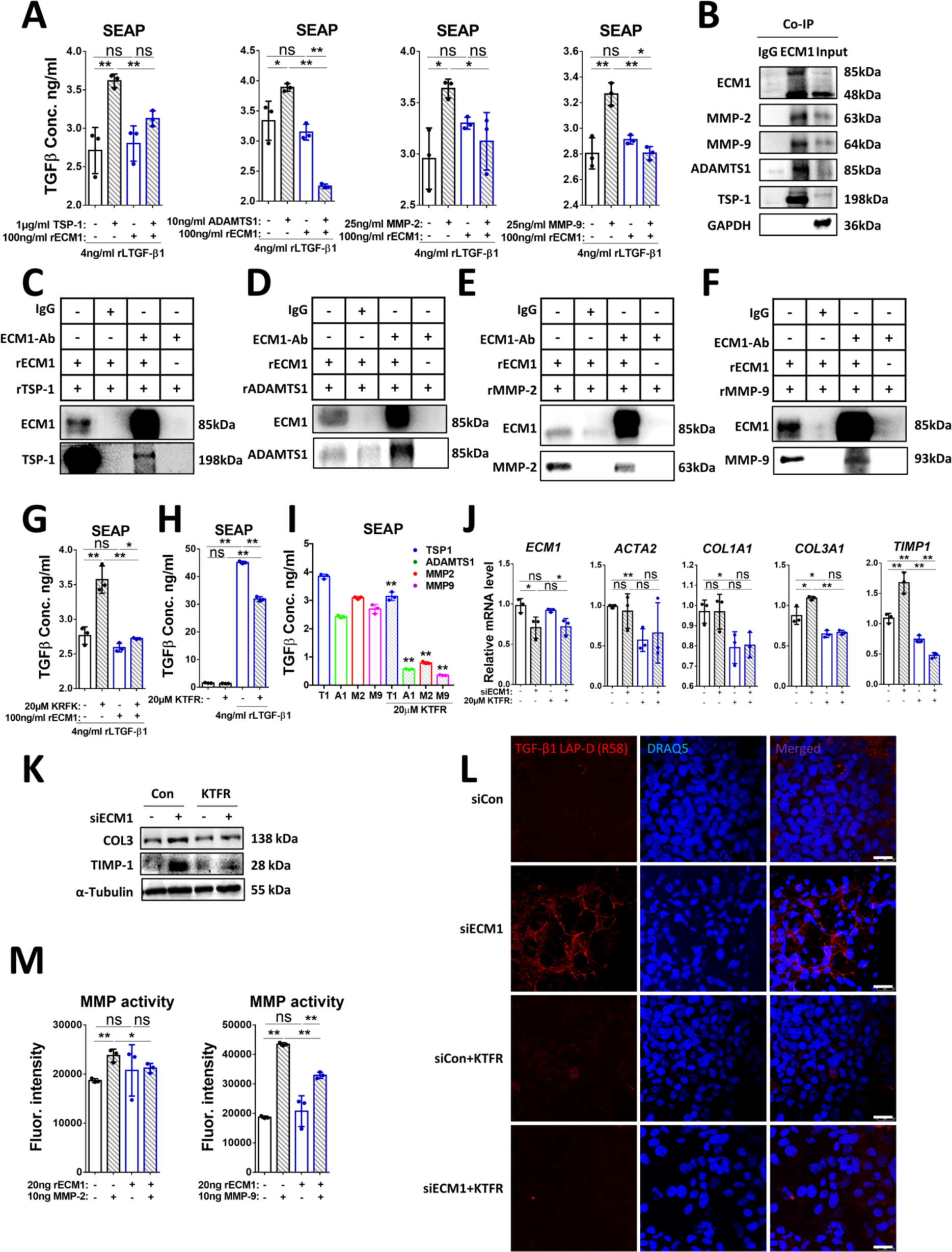
ECM1 blocks TSP-1- and ADAMTS1-mediated LTGF-β1 activation via an interaction with their respective KRFK/KTFR amino acid motifs and inhibits the proteolytic activity of MMP-2 and MMP-9. **(A)** The concentration of active TGF-β (ng/ml) measured by SEAP activity assay from conditioned MFB-F11 SN incubated with/without rLTGF-β1, rECM1 and/or LTGF-β1 activators. **(B)** Immunoprecipitation studies of ECM1 with MMP-2, MMP-9, ADAMTS1, or TSP-1 in LX-2 HSCs cultured for 72h. An anti-human ECM1 antibody was used for immunoprecipitation. MMP-2, MMP-9, ADAMTS1, and TSP-1 antibodies were used for Western blotting. **(C-F)** *In vitro* pull-down assays of ECM1 and TSP-1, ADAMTS1, MMP-2, or MMP-9. Recombinant proteins of ECM1, TSP-1, ADAMTS1, MMP-2, or MMP-9 were incubated with protein A/G agarose. IgG or anti-human ECM1 antibody was used for immunoprecipitation. ECM1, MMP-2, MMP-9, ADAMTS1, and TSP-1 antibodies were used for Western blotting. **(G)** The concentration of active TGF-β (ng/ml) measured by SEAP activity assay from conditioned MFB-F11 SN incubated with/without rLTGF-β1, rECM1 and/or KRFK peptide for 24h. **(H)** The concentration of active TGF-β (ng/ml) measured by SEAP activity assay from conditioned MFB-F11 SN incubated with/without rLTGF-β1 and/or KTFR peptide for 24h. **(I)** The concentration of active TGF-β (ng/ml) measured by SEAP activity assay from conditioned MFB-F11 SN incubated with rLTGF-β1 and the four mediators respectively combined the incubation with/without KRFK peptide for 24h. **(J)** Relative mRNA levels of *ACTA2*, *COL1A1*, *COL3A1*, and *TIMP1* in LX-2 HSCs transfected with siCon or si*ECM1*, incubated with/without KTFR for 48h. **(K)** Protein expression of COL3 and TIMP1 in LX-2 HSCs with the treatment as shown in the figure. **(L)** IF staining of TGF-β1 LAP-D (R58) in LX-2 HSCs with the treatment as shown in the figure. Scale bar, 25 µm. **(M)** MMP activity assay showing the enzymatic activity of MMP-2 and MMP-9 with/without rECM1 incubation. MMP Green Substrate was used as a control. For RT-qPCR, human PPIA was used as an endogenous control. *P*-values were calculated by unpaired Student’s t test. Bars represent the mean ± SD. *, *P*<0.05; **, *P*<0.01.

It is known that TSP-1 and ADAMTS1 activate LTGF-β1 through an interaction of their respective KRFK [32] or KTFR [33] amino acid motifs. Both motifs are said to induce a conformational change in the LTGF-β1 molecule to release the active TGF-β1 ligand from its complex without the need for additional proteolysis. Further, there is data indicating TSP1’s KRFK peptide in being sufficient to activate LTGF-β1, while the KTFR peptide within the full ADAMTS1 protein is necessary, but by itself not sufficient for LTGF-β1 activation. In fact, the KTFR peptide displays a competitive inhibition of ADAMTS1-mediated LTGF-β1 activation when used in a recombinant form [26, 33]. To identify how ECM1 functionally integrates into these seemingly opposing effects of the KRFK and KTFR peptides on LTGF-β1 activation, we tested chemically synthesized KRFK and KTFR peptides in two distinct contexts. An inert KQFK peptide was used as a negative control. MFB-F11 cells were incubated with rLTGF-β1 followed by treatment of rECM1 and/or KRFK peptide to demonstrate that rECM1 caused marked inhibition of KRFK peptide-induced effects on the TGF-β reporter assay activity **(Figure 4G)**. Due to different properties of the ADAMTS1-KTFR motif, MFB-F11 cells were treated with rLTGF-β1 and/or KTFR for 24h to show that KTFR inhibits LTGF-β1 auto-activation. This could be demonstrated by the SEAP assay **(Figure 4H)**. Furthermore, KTFR pre-treatment inhibited TSP-1-, ADAMTS1-, MMP-2-, and MMP-9-mediated rLTGF-β1 activation **(Figure 4I)**. In addition, treatment of LX-2 HSCs with KTFR reduced siECM1-induced active TGF-β1 levels, as evident from mRNA expression of *ACTA2*, *COL1A1*, *COL3A1*, and *TIMP1* **(Figure 4J)**, protein levels of COL3A1 and TIMP1 **(Figure 4K)**, and IF staining of TGF-β1 LAP-D R58 **(Figure 4L)**.

MMPs mediate LTGF-β1 activation by proteolytic cleavage of LAP [27]. To investigate whether ECM1 affects the proteolytic activity of MMP-2 or MMP-9, we performed fluorogenic MMP activity assays. Incubation of MMP-2 or MMP-9 with rECM1 at a ratio of 1:2 (20ng MMP-2 or -9 and 40ng rECM1) inhibited the proteolytic activity of MMP-2 and MMP-9 **(Figure 4M)**.

In summary, we show that the abrogative effect of ECM1 on LTGF-β1 activation by TSP-1 or ADAMTS1 integrates at the KRFK or KTFR amino acid sequence. Additionally, ECM1 binds to mature MMP-2 and MMP-9 and thereby inhibits their proteolytic activity, thus reducing LAP degradation and LTGF-β1 activation.

### Overexpression of ECM1 inhibits KRFK-induced LTGF-β1 activation in murine livers

Since a KRFK peptide challenge phenocopied TSP-1’s function regarding LTGF-β1 activation, we expected that treatment of mice with the KRFK peptide would lead to higher levels of active TGF-β1 and increased fibrogenesis. To test whether ECM1 can prevent KRFK-induced LTGF-β1 activation *in vivo* and thus rescue any resulting liver injury, we treated 8-weeks-old WT mice with control or AAV8-ECM1 for 7 days prior to daily intraperitoneal injections of 100 µg KRFK or KQFK (negative control) peptide for 14 consecutive days **(Figure 5A)**. The KRFK injections caused a significant upregulation of HSC-specific fibrosis markers such as *Acta2*/α-SMA **(Figure 5B-D)** and collagen types *Col1α1* and *Col3α1* **(Figure 5B)** all of which were also upregulated in 2- and 5-weeks-old *Ecm1*-KO mice **(Figure 1I)**. Further, IHC staining of TGF-β1 LAP-D R58, H&E, and picrosirius red (PSR) staining demonstrated that KRFK peptide treatment caused significant LTGF-β1 activation and hepatocyte injury as displayed by cleared-out cytoplasm, and pericentral liver fibrosis **(Figure 5E)**. Cell apoptosis was assessed by IHC staining of cleaved caspase 3 to further investigate the type of KRFK-mediated liver injuries, however, we did not detect increased hepatocyte apoptosis compared to the control group (**Figure S5A**). This suggests that KRFK treatment is sufficient to induce mild fibrosis, however is not sufficient for hepatocyte apoptosis levels. In turn, prior AAV8-mediated overexpression of ECM1 prevented the pro-fibrogenic changes induced by the KRFK peptide, as evident from reduced levels of *Acta2*/α-SMA **(Figure 5B-D)**, decreased mRNA expression of fibrillar collagens type *Col1α1* and *Col3α1* **(Figure 5B)**, and less detection of TGF-β1 LAP-D R58 **(Figure 5E)**. Further, AAV8-ECM1 pre-treated mice presented with less evidence of liver injury in H&E staining and fewer areas of fibrosis on PSR staining. Taken together, ECM1 overexpression effectively prevents liver injury and fibrosis caused by KRFK-mediated activation of LTGF-β1, suggesting a potential benefit in TSP-1-driven disease settings [34].

**Figure 5.**
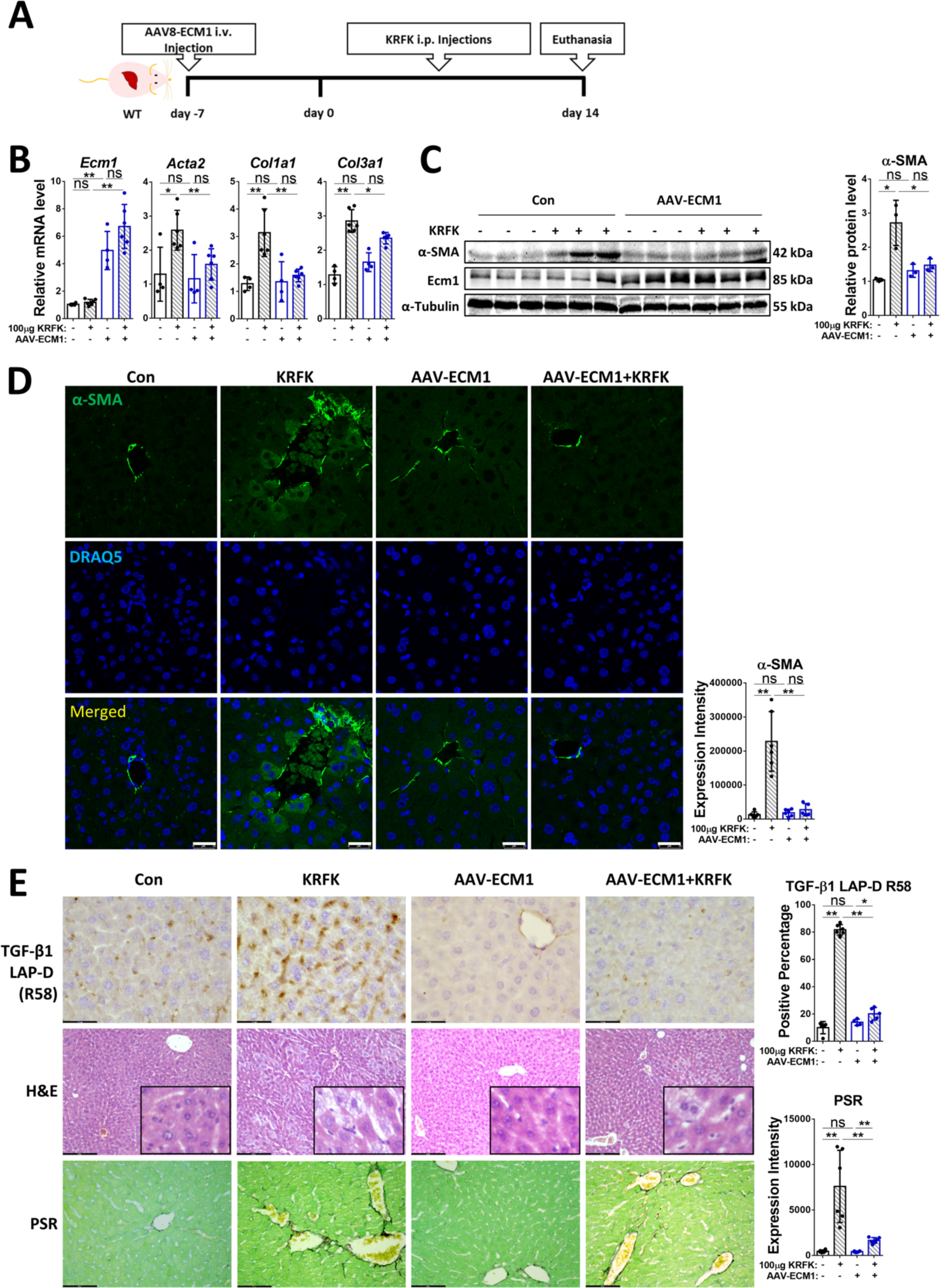
ECM1 overexpression in the murine liver prevents KRFK peptide-induced LTGF-β1 activation and subsequent hepatic fibrosis in WT mice. **(A)** Schematic illustration of *in vivo* experiment with AAV-ECM1 (i.v.) and KRFK peptide (i.p.) injections of WT mice. **(B)** Relative RT-qPCR mRNA levels of *Ecm1*, *Acta2*, *Col1a1*, and *Col3a1* in liver tissue from WT mice treated w/o AAV-ECM1, plus control or KRFK peptide injection daily, for 14 consecutive days. **(C)** Protein levels of α-SMA in liver tissue from WT mice treated with control or AAV-ECM1 followed by control or KRFK peptide. **(D)** Representative images of IF stainings for α-SMA in liver tissue from WT mice treated with control or AAV-ECM1 followed by control or KRFK peptide. Scale bar, 25 µm. **(E)** Representative images of IHC stainings for TGF-β1 LAP-D R58, H&E, and PSR staining in liver tissue from WT mice treated with control or AAV-ECM1 followed by control or KRFK peptide. Scale bar, 43.5, 174, 87 µm, respectively. For RT-qPCR, mouse *Ppia* was used as an endogenous control. For Western blotting, α-Tubulin was used as a loading control. P-values were calculated by unpaired Student’s t test. Bars represent the mean ± SD. *, *P*<0.05; **, *P*<0.01.

### KTFR reverses *Ecm1*-KO-induced liver injury in mice

As KTFR suppressed LTGF-β1 activation *in vitro* **(Figure 4H-L)**, we injected 100 µg KTFR peptide into 8-weeks-old *Ecm1*-KO mice for 14 consecutive days to test if we could ameliorate the LTGF-β1 activation causing progressive liver injury in *Ecm1*-KO mice **(Figure 6A)**. A KQFK peptide was used as a negative control. Surprisingly, KTFR treatment was able to revert the *Ecm1*-KO phenotype. RT-qPCR and immunoblotting data showed that fibrosis markers in *Ecm1*-KO livers were attenuated by KTFR peptide injections, including *Col1α1*, *Col3α1*, *Timp1*, and *Acta2*/α-SMA **(Figure 6B, C)**. IF or IHC staining of α-SMA and TGF-β1 LAP-D (R58) further confirmed the inhibitory effect of KTFR on *Ecm1*-KO-mediated HSC and LTGF-β1 activation **(Figure 6D, E)**. H&E and PSR stainings exhibited perisinusoidal fibrosis in *Ecm1*-KO mice as shown by extracellular matrix deposition being attenuated by KTFR peptide injection **(Figure 6E)**. In addition, 10-week-old *Ecm1*-KO mice exhibited cell apoptosis in a few hepatocytes, characterized by nuclear expression of cleaved caspase 3 and cell shrinkage compared to WT mice, which was alleviated by KTFR treatment (**Figure S5B**, indicated by arrows).

**Figure 6.**
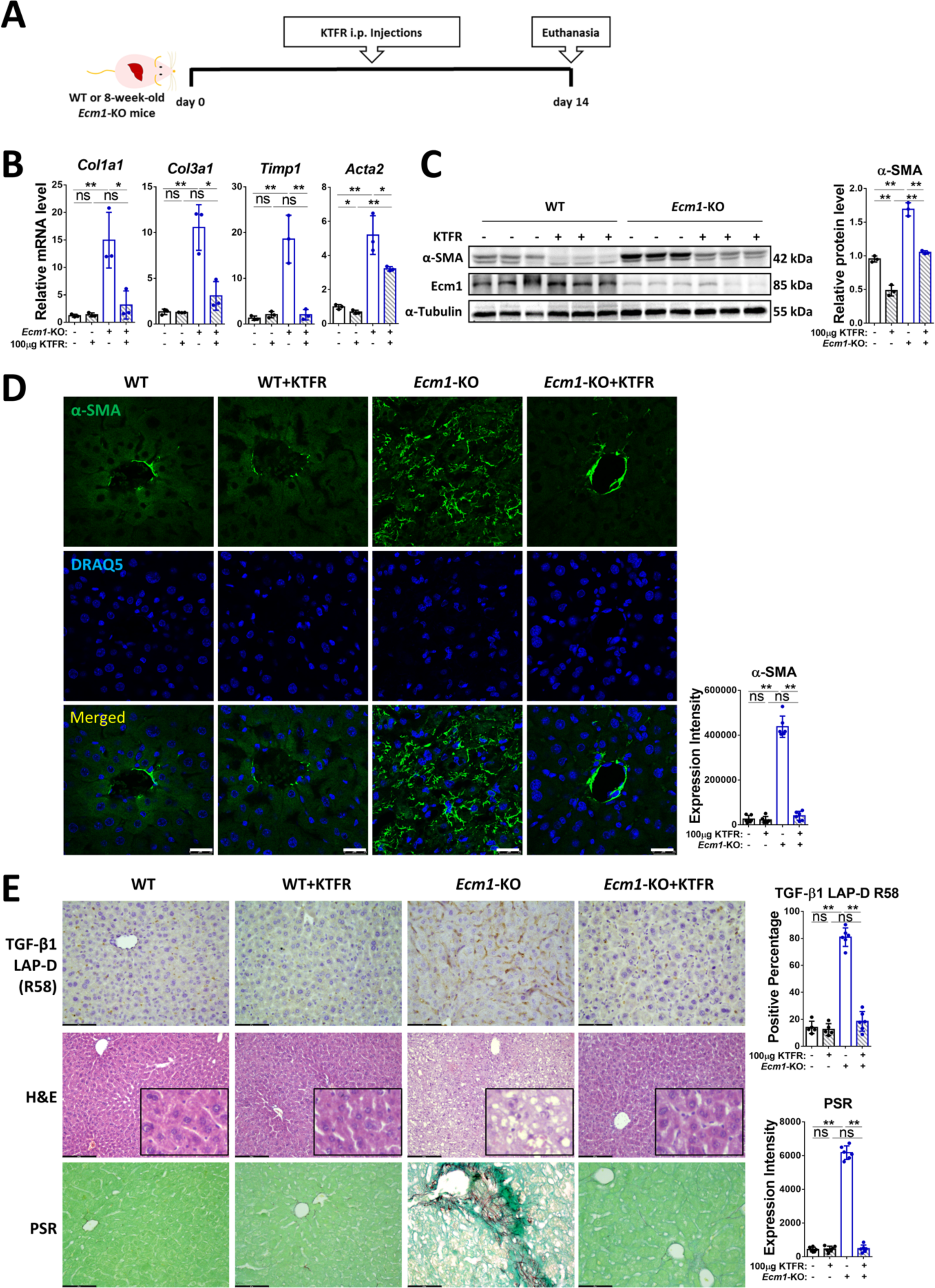
KTFR peptide treatment rescues liver injury in *Ecm1*-KO mice. **(A)** Schematic illustration of *in vivo* experiments with KTFR peptide injections (i.p.) into WT and *Ecm1*-KO mice. **(B)** Relative mRNA levels of *Col1a1*, *Col3a1*, *Timp1*, and *Acta2* in liver tissue from WT and *Ecm1*-KO mice injected with control or KTFR peptide daily, for 14 consecutive days. **(C)** Protein levels of α-SMA in liver tissue from WT and *Ecm1*-KO mice injected with control or KTFR peptide. **(D)** Representative images of IF stainings of α-SMA in liver tissue from WT and *Ecm1*-KO mice injected with control or KTFR peptide. Scale bar, 25 µm. **(E)** Representative images of IHC stainings for TGF-β1 LAP-D R58, H&E, and PSR staining in liver tissue from WT and *Ecm1*-KO mice injected with control or KTFR peptide. Scale bar, 43.5, 174, 87 µm, respectively. For RT-qPCR, mouse *Ppia* was used as an endogenous control. For Western blotting, α-Tubulin was used as a loading control. *P*-values were calculated by unpaired Student’s t test. Bars represent the mean ± SD. *, *P*<0.05; **, *P*<0.01.

*In vivo*, KTFR peptide exerts an effect similar to that of ECM1 and reverts the liver injury suffered by mice upon *Ecm1-*KO at 8 weeks of age and could serve to restore ECM1’s functions where lost.

### KTFR alleviates liver injury in *Fxr*-KO mice

Farnesoid X receptor (FXR, NR1H4) plays fundamental roles in maintaining bile acid homeostaisis, modulating metabolism, and reducing inflammation [35]. *Fxr*-KO mice are characterized by cholestatic liver injury, HSC activation, fibrosis, and subsequcently progression to HCC [35, 36]. We observed that the expression of *Ecm1* was significantly decreased in *Fxr*-KO mice while TGF-β1 mRNA expression remained unaffected (**Figure S6A**), which suggested that fibrosis in *Fxr*-KO mice was primarily due to LTGF-β1 activation. In consequence, *Fxr*-KO mouse model may represent an ideal platform for further validation and asessment of the anti-fibrotic potential of the KTFR peptide. Therefore, 100 µg KTFR or KQFK (negative control) peptide were injected into 8-week-old *Fxr*-KO mice for 14 consecutive days (**Figure 7A**). RT-qPCR and immunoblotting analyses demonstrated that the increased fibrosis markers in *Fxr*-KO livers, including *Col1α1*, *Col3α1*, *Timp1*, and *Acta2*, were rescued following KTFR peptide treatment **(Figure 7B, C)**. IF or IHC staining of α-SMA and TGF-β1 LAP-D (R58) further confirmed the inhibitory effect of KTFR on HSC and LTGF-β1 activation in *Fxr*-KO mice **(Figure 7D, E)**. H&E and PSR stainings revealed pericentral fibrosis in *Fxr*-KO mice, characterized by extracellular matrix deposition around the central veins, which was attenuated by KTFR peptide injection **(Figure 7E)**. Similar to *Ecm1*-KO mice, mild hepatocyte apoptosis induced by *Fxr*-KO was prevented by KTFR treatment, as assessed by IHC staining of cleaved caspase 3 (**Figure S6B**, indicated by arrows). These results confirmed the anti-fibrotic effect of KTFR in *Fxr*-KO mice.

**Figure 7.**
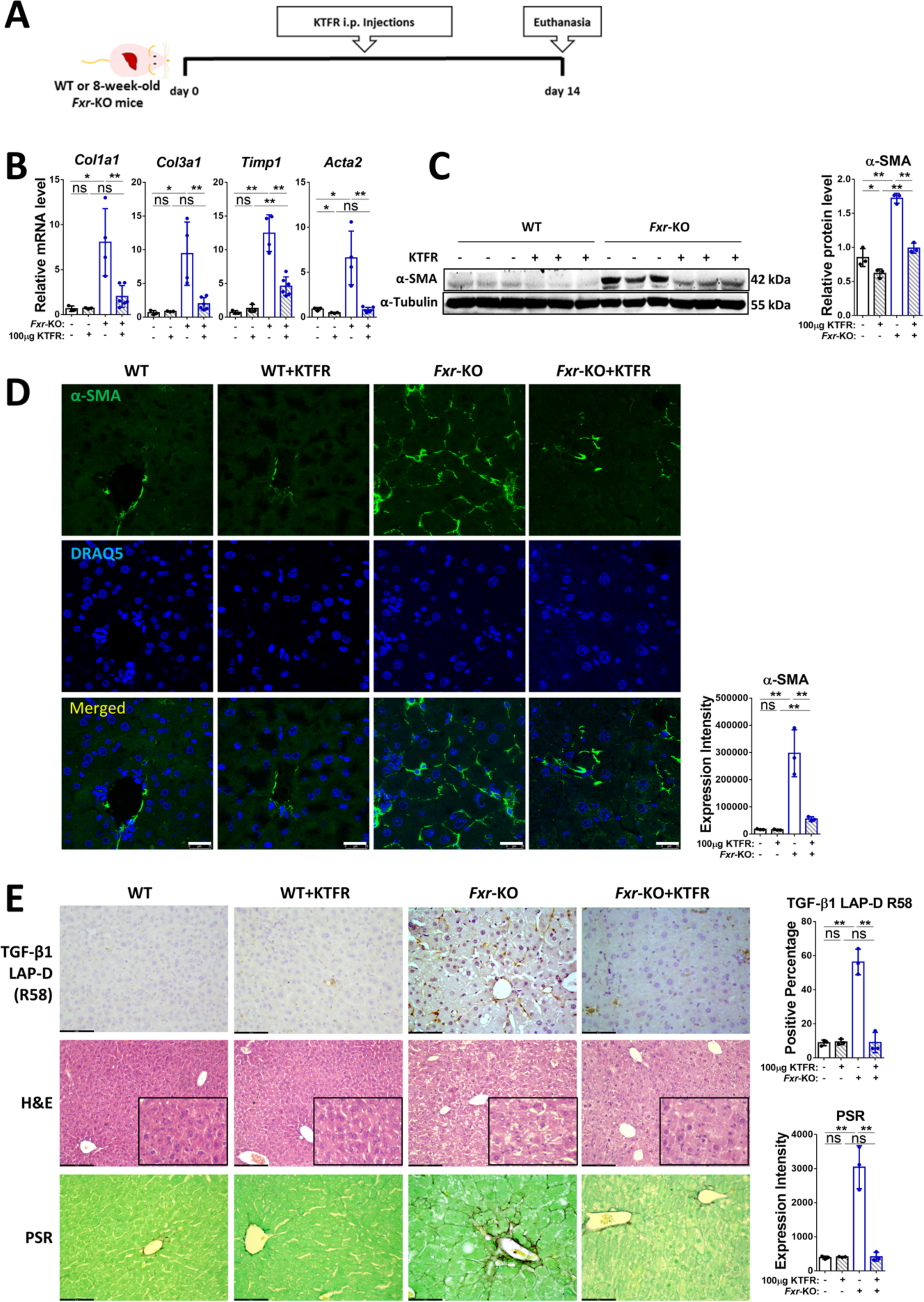
KTFR peptide treatment rescues liver injury in *Fxr*-KO mice. **(A)** Schematic illustration of *in vivo* experiments with KTFR peptide injections (i.p.) into WT and *Fxr*-KO mice. **(B)** Relative mRNA levels of *Col1a1*, *Col3a1*, *Timp1*, and *Acta2* in liver tissue from WT and *Fxr*-KO mice injected with control or KTFR peptide for 14 consecutive days. **(C)** Protein levels of α-SMA in liver tissue from WT and *Fxr*-KO mice injected with control or KTFR peptide. **(D)** Representative images of IF stainings of α-SMA in liver tissue from WT and *Fxr*-KO mice injected with control or KTFR peptide. Scale bar, 25 µm. **(E)** Representative images of IHC stainings for TGF-β1 LAP-D R58, H&E, and PSR staining in liver tissue from WT and *Fxr*-KO mice injected with control or KTFR peptide. Scale bar, 43.5, 174, 87 µm, respectively. For RT-qPCR, mouse *Ppia* was used as an endogenous control. For Western blotting, α-Tubulin was used as a loading control. *P*-values were calculated by unpaired Student’s t test. Bars represent the mean ± SD. *, *P*<0.05; **, *P*<0.01.

### ECM1 loss correlates with upregulated mediators of LTGF-β1 activation in CLD

To investigate the relationship between ECM1 downregulation and LTGF-β1 activation in patients, we performed RNAscope, IF or IHC staining for ECM1, TGF-β1 LAP-D (R58), TSP-1, ADAMTS1, MMP-2 and MMP-9 in paraffin-embedded tissue samples from patients suffering from either F1-F2 liver fibrosis (n=6) or ALD-/HBV-induced liver cirrhosis (n=15) and compared those to relatively healthy individuals (n=5). In parallel with the progressive decrease of ECM1 expression on RNA and protein level **(Figure 8A, B)**, TGF-β1 LAP-D (R58) staining intensity increased from control to F1-2 stage fibrosis and on to cirrhosis **(Figure 8C)** indicating a significant and inverse correlation. Interestingly, expression of the tested LTGF-β1 activators, TSP-1, ADAMTS1, MMP-2 and MMP-9 increased consistently with progressing liver fibrosis **(Figure 8C)**. This suggests that in parallel with ECM1 downregulation, the bioavailability of the mediators increase as well in order to provide active TGF-β1 for surrounding liver tissue in its response to damage. This was further confirmed across different etiologies of liver injury by analysing the GEO DataSets (GSE49541) that comprises RNAseq data of liver tissue from NAFLD patients with mild (n=40) or advanced (n=32) fibrosis [37]. ECM1 expression decreased with disease severity, while mRNA levels of LTGF-β1 activators increased including that of *TSP-1*, *TSP-2*, *ADAMTS1*, *ADAMTS2*, *MMP-2*, *MMP-9*, *ITGAV*, and *ITGB1*. As this would lead to enhanced availability of active TGF-β1, downstream activation of HSCs can be expected, which was supported by increased expression levels of fibrotic collagens, such as *COL1A1*, *COL1A2*, *COL3A1*, and *COL4A1* in cohorts with more advanced fibrosis **(Figure 8D)**. Additionally, we analyzed human single cell (sc)RNAseq (GSE174748) of liver tissue from normal and NAFLD cirrhotic samples [7]. After quality filtering, 30,918 cells, with an average sequencing depth of 2,000 genes per cell, were subjected to further analysis. Unsupervised clustering classified these cells into 14 different clusters **(Figure S7A)**, and Cluster 10 was defined as HSCs based on the expression of HSC-specific marker genes **(Figure S7B)**. Uniform manifold approximation and projection (UMAP) visualization showed that HSCs could be divided into two clusters: quiescent HSCs (qHSCs) and active HSCs (aHSCs) **( Figure S7C, 8E)**. With *ECM1* was abundant in quiescent HSCs, LTGF-β1 activators (*TSP-1*, *ADAMTS1*, *MMP-2*, *MMP-9*), and fibrotic markers (*ACTA2*, *COL1A1*) were highly enriched in activated HSCs (**Figure 8F, G**). In consequence, we hypothesize that the sequence of events in CLD expression starts with ECM1 downregulation and upregulation of LTGF-β1 activators, followed by LTGF-β1 and HSC activation and finally, increased extracellular matrix production and deposition **(Figure 9A, B)**.

**Figure 8.**
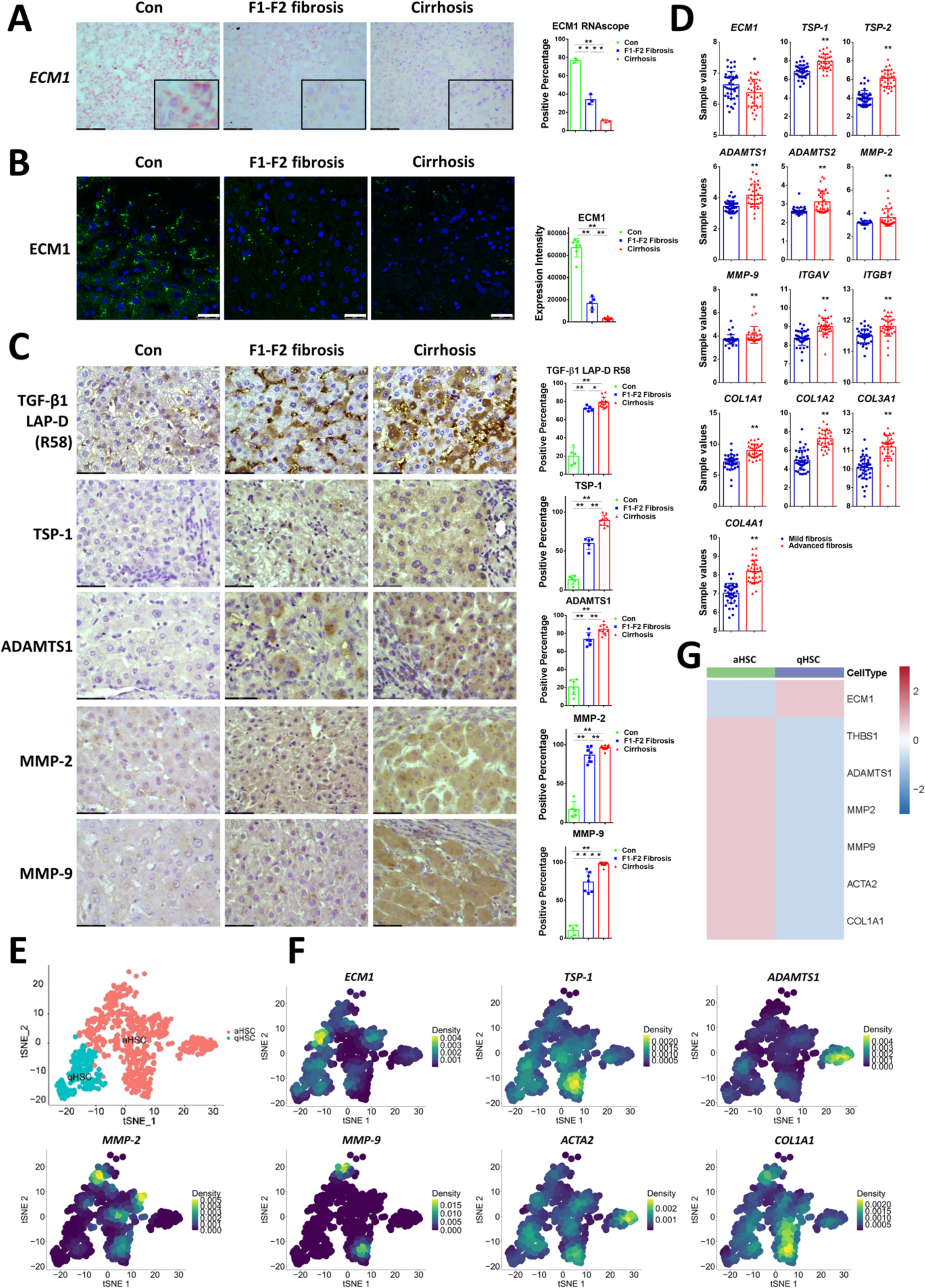
Decreased ECM1 and increased mediators of L-TGF-β1 activation are associated with progressing fibrosis in CLD patients. **(A)** Representative *ECM1* RNAscope data in liver tissue from control, F1-F2 fibrosis, and cirrhosis liver tissue. Scale bar, 87 µm. **(B)** Representative images of IF stainings for ECM1 in control, F1-F2 fibrosis, and cirrhosis liver tissue. DRAQ5 was used for nuclear staining. Scale bar, 25 µm. **(C)** Representative images of IHC staining for TGF-β1 LAP-D R58, TSP-1, ADAMTS1, MMP-2, and MMP-9 in control, F1-F2 fibrosis, and cirrhosis liver tissue. Scale bar, 43.5 µm. **(D)** mRNA expression of *ECM1*, LTGF-β1 activators, and fibrotic collagens in liver tissue from patients with F1-2 fibrosis or cirrhosis, extracted from GEO DataSets GSE49541. **(E)** UMAP visualization of isolated hepatic stellate cells from GSE174748. **(F, G)** Density plots and heatmaps show the relative expression of target genes in scRNAseq of HSC subsets from GSE174748. Quantification of stainings was performed using Image J (National Institute of Health, Bethesda, MD). *P*-values were calculated by unpaired Student’s t test. Bars represent the mean ± SD. *, *P*<0.05; **, *P*<0.01.

**Figure 9.**
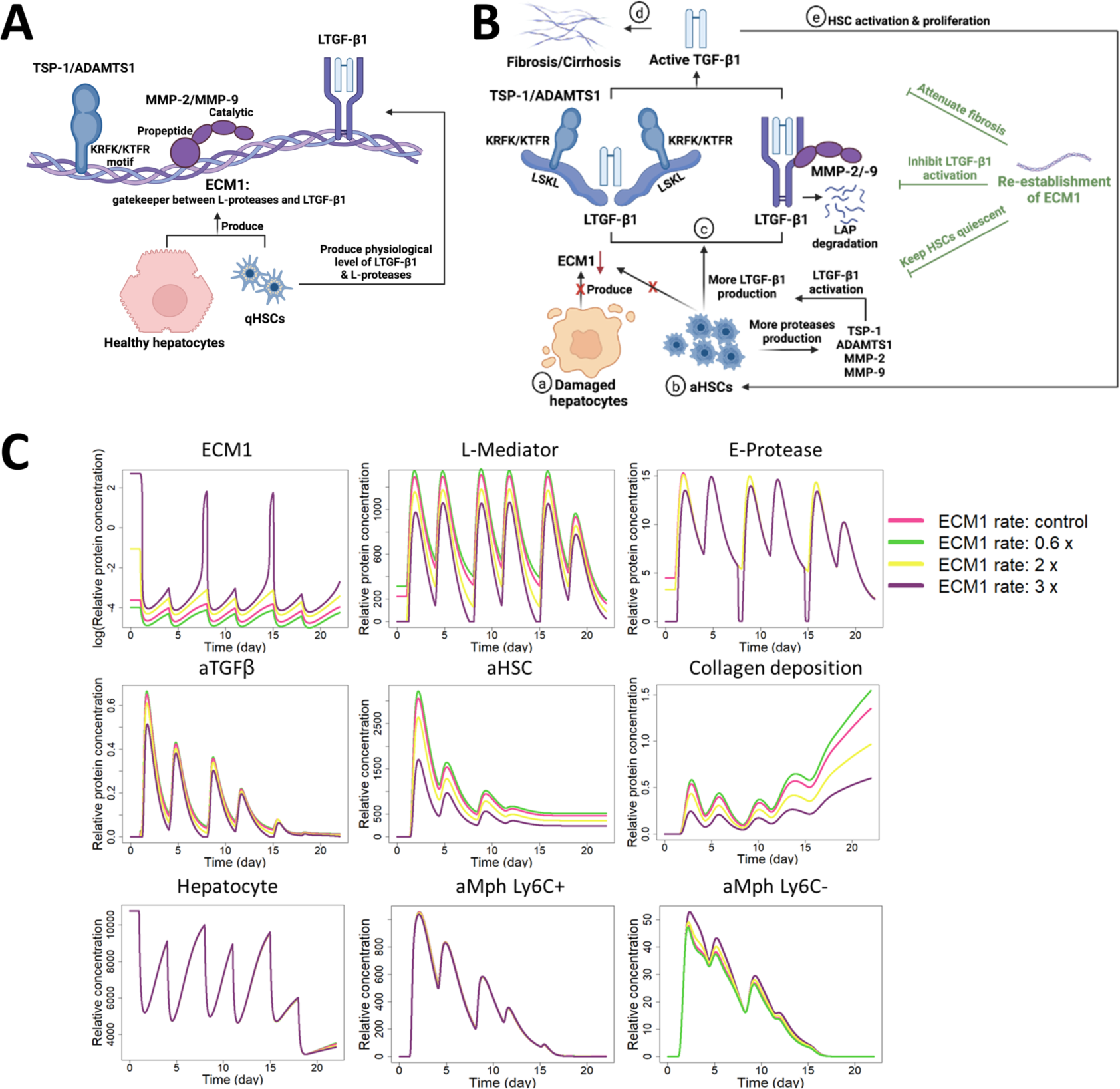
ECM1’s role in maintaining liver tissue homeostasis in healthy livers and its dysregulation during chronic liver disease. **(A)** In healthy livers, ECM1 from healthy hepatocytes and quiescent HSCs prevents excessive LTGF-β1 activation by TSP-1 and ADAMTS1 through interacting with their respective KRFK and KTFR motifs and binding to MMP-2 an MMP-9, limiting their enzymatic activity thus keeping the TGF-β1 inactive within its ECM-deposited latent complex (LTGF-β1). **(B)** During CLD, (a) damaged hepatocytes fail to maintain ECM1 expression and production which results in (b) increased and detrimental LTGF-β1 activation via (c) conformational changes in the LAP molecule caused by TSP-1 and ADAMTS1 or via direct proteolysis of the LAP molecule as caused by MMP-2 and MMP-9. Increased levels of active TGF-β1 promote (d) the accumulation of excessive extracellular matrix and cause (e) HSC activation, thus resulting in (progressive) hepatic fibrosis. Intriguingly, ectopic ECM1 expression (rECM1/v-ECM1) inhibits TSP-1, ADAMTS1, MMP-2 and MMP-9-mediated LTGF-β1 activation and attenuates subsequent hepatic fibrosis (Right in green). **(C)** Simulation of chronic liver injury. Simulation of 4 doses of *CCl*_*_ (at day 1, 4, 8, 11) for 15 days under four different ECM1 production rates: the control *k*^-.^, *k*^--^, 0.6-fold of the control [[umath]] 0.6 × *k*^-.^, 0.6 × *k*^--^, 2-fold of the control 2 × *k*^-.^, 2 × *k*^—^[[umath]], and 3-fold of the control 3 × *k*^-.^, 3 × *k*^--^. The first dose of *CCl*_*_is injected at day 1 when the system is in a healthy steady state. Shown are the relative concentrations of ECM1, L-Mediator and E-Protease (first row), active TGF-β1 (aTGF-β1), activated HSC (aHSC), collagen deposition (second row), hepatocytes, Ly6C+ and Ly6C-macrophages (aMph) (third row) as a function of time for four different ECM1 production rates.

### Compartment modeling confirms consistency of effects of ECM1 on the liver fibrosis interaction network

In order to verify the experimentally observed behavior of ECM1, we established a computational model that represents the key network of cellular phenotypes and their hypothesized interactions **(Figure S8)**. To simulate the process of fibrogenesis, we ran the simulation for 21 days which represents 6 injections of *CCl*_*_(at days 1, 4, 8, 11, 15 and 18, following the experimental administration scheme) under four different ECM1 production rates as shown in Figure 9C. The first dose of *CCl*_*_was injected at day 1, when the equilibrium was reached. The overall temporal dynamics of the displayed network components well reproduced the known or expected behavior of chronic liver injury. When the production rate of ECM1 was increased in the simulation, the amount of E-Protease (degrades ECM1 in our model), L-Mediator (activates LTGF-β1 in our model) and active TGF-β1 decreased. The quantity of aHSC and fibrotic collagen dropped as the production rate of ECM1 increased. The mass of hepatocytes and macrophages in the compartment, however, was not influenced, even though the small dispersion of the curves for hepatocytes at 20 days indicates that in case of even higher fibrotic collagen concentrations, the hepatocyte mass may be affected by ECM1 production **(Figure 9C)**. In summary, when incorporating the role of ECM1 as delineated in our experiments, the compartment model accurately describes development and progression of hepatic fibrosis following chronic liver injury. The reduced levels of ECM1 lead to increased production of fibrotic matrix. This supports the experimental conclusion that ECM1 represents a suitable therapeutic target to reverse hepatic fibrosis.

## Discussion

The present study shows that ECM1 inhibits LTGF-β1 activation mediated by TSP-1, ADAMTS1, MMP-2 and MMP-9 and thus attenuates hepatic fibrosis **(Figure 9A, B)**. TSP-1 and ADAMTS1 cause a conformational change within LAP of the LTGF-β1 complex, through their respective KRFK or KTFR motifs binding to LAP’s LSKL sequence, thereby releasing the active cytokine [32, 38, 39]. Interestingly, while the KRFK peptide can independently activate LTGF-β1**(Figure S9A, B)**, KTFR exposure requires collaboration with another motif within ADAMTS1 to release the active TGF-β1 **(Figure S9C, D)**. Thus, the presence of a KTFR peptide alone suffices to maintain the integrity of LTGF-β1, not allowing its activation. In contrast, MMP-2 and MMP-9 act via proteolytical cleavage of LAP, resulting in LTGF-β1 activation [27]. It has been shown that the C-terminal tandem repeat 2 (amino acids 236-361) of ECM1 binds to MMP-9 thus reducing its activity [40]. Whether this C-terminal repeat or any other sequence of ECM1 interacts with MMP-2 needs further investigation.

To further verify our findings in liver tissue of patients suffering from CLD, we show that with ECM1 downregulation and LTGF-β1 activation, there is a significantly increased expression of LTGF-β1 activating mediators, including TSP-1, ADAMTS1, MMP-2, and MMP-9 which correlate with the progressing fibrosis **(Figure 8A-C)**. It is known that activated HSCs and injured hepatocytes are the major source for TSP-1 [41], ADAMTS1 [42], as well as MMP-2 and MMP-9 [43] in fibrotic liver disease, indicating that expression levels of LTGF-β1 activators are useful parameters to predict liver damage. It remains a challenge to directly investigate the activity of TSP-1, ADAMTS1, MMP-2, and MMP-9 in patient liver tissue, but we were able to show their activity indirectly by staining for TGF-β1 LAP-D R58.

To delve deeper into the impact of these mediators on TGF-β1 activation, we examined the mRNA expression of TGF-β1 in LX-2 HSCs treated with control or αvβ6 integrin/TSP-1/ADAMTS1/MMP-2/MMP-9 and found an induction of TGF-β1 mRNA expression **(Figure S10A)**. Whether it stems from TGF-β1’s positive and self-sustained feedback loop on its own expression or from LTGF-β1 activators facilitating TGF-β1 transcription remains inadequately explored and warrants further study. We hypothesize that reverting the loss of ECM1 might disrupt the observed self-perpetuating circle of TGF-β1 activation and expression. Reversely, we also investigated whether ECM1 affected the expression of its activators. ECM1 overexpression inhibited the mRNA expression of αvβ6 integrin, TSP-1, ADAMTS1, MMP-2, and MMP-9, while rECM1 did not significantly reduce their expressions (**Figure S10B, C**). These findings in combination with our interaction studies and SEAP assays (**Figure 3, 4**) suggest that ECM1 does indeed inhibit mediator-induced LTGF-β1 activation, but it may also attenuate liver fibrosis by preventing their expression. It is currently not clear how ECM1 could control the expression of TSP-1/ADAMTS1/MMP-2/MMP-9 beyond TGF-β1 bioavailability. Additionally, we investigated the effects of ECM1 on HSC behavior by ECM1 overexpression or knockdown in LX-2 cells. ECM1 overexpression significantly inhibited cell proliferation (**Figure S11A-C**) and migration (**Figure S11D**), while ECM1 knockdown did not affect proliferation (**Figure S11E-G**) but appeared to promote cell migration (**Figure S11H**). As LX-2 cells are activated HSCs with a comparatively low level of ECM1 expression versus quiescent HSCs, further analyses with ECM1 knockdown should be conducted in primary HSCs to provide more relevant insights.

ECM1 is predominantly expressed by hepatocytes and to a lesser extent by quiescent HSCs, which secrete it into the liver’s extracellular matrix, to protect the matrix-deposited LTGF-β1 from activation by the mediators and subsequent fibrogenic signaling. In addition to HSCs and TGF-β1, CLD progression involves a plethora of cell types and signals. Therefore, we used computational modelling to further investigate the hepatoprotective effect of ECM1 **(Figure 9C)**. The simulated temporal dynamics of the proposed *in silico* model predict that indeed ECM1 availability might be a limiting factor that can be overcome to attenuate injury in CLD. Extending this network in the future and resolving it in space and time, as recently demonstrated for liver regeneration [44], will inform a future multi-scale model, or a digital liver twin. Precise *in silico* modelling will allow researchers to predict the impact of the up- or down-regulation of each element on all other network components and could lead to a better understanding of the entire disease process, and help us anticipate the possible success of future therapeutic interventions.

Reliable and safe delivery to the human liver is essential for antifibrotic treatment. It may be possible to employ recombinant proteins or phenocopying peptides [45], viral vectors such as adeno-associated viruses (AAV), which have already been employed in gene therapy [46], bacterial delivery [47] methods to restore ECM1 expression or function in diseased livers.

Short synthetic peptides are a promising strategy, as they have been successfully employed in the treatment of multiple sclerosis and NASH [48, 49]. Therefore, a peptide sequence of the ECM1 protein that confers its function on LTGF-β1 bioavailability and prevents it from excessive activation or a direct application of the KFTR peptide as shown experimentally *in vivo* might provide a liver-specific, anti-fibrotic intervention, which are still not available at the present time [49, 50].

## Materials and Methods

Human samples, mouse experiments, chemical reagents, primers, antibodies used in this study, and further detailed information on the methods are presented in the supplementary materials.

## Author contributions

Conceptualization: SD, SW

Methodology: FL, SW, YL, JZ, DD, SM, WF, ZCN, YY, CH, SH

Investigation: FL, SW, YL, JZ, SM, ZCN, WF, SW, YY, HL, CS, DD

Visualization: FL, SW, JZ, DD, YY, CH

Funding acquisition: SD

Project administration: SD, SW, FL

Supervision: SW, SD

Writing - Original: FL, SW, SD

Writing - review: FL, SW, SD, JZ, DD, PTD, ZCN, RL, HW, ME, NJT, BS, HD, CG

SW and SD acted as guarantors.

## Acknowledgments

We acknowledge the support of the LIMa Live Cell Imaging at Microscopy Core Facility Platform Mannheim (CFPM).

## Abbreviations

aHSC: activated HSCs
aMph: Ly6C+ and Ly6C-macrophages
aTGF-β: active TGF-β
AAV: adeno associated virus
ADAMTS1: A disintegrin and metalloproteinase with thrombospondin motifs 1
ASH: Alcoholic steatohepatitis
*CCl_4_*: carbon tetrachloride
CLD: Chronic liver disease
ECM1: Extracellular matrix protein 1
FXR: Farnesoid X recptor
HCC: Hepatocellular carcinoma
HSC: Hepatic stellate cell
IF: Immunofluorescence staining
IHC: Immunohistochemistry staining
i.v.: intravenous
i.p.: intraperitoneal
KO: knockout
KQFK: Lysine-Glutamine-Phenylalanine-Lysine
KRFK: Lysine-Arginine-Phenylalanine-Lysine
KTFR: Lysine-Threonine-Phenylalanine-Arginine
LAP: Latency associated peptide
LTGF-β (1): Latent transforming growth factor-β (1)
MMP-2: matrix metalloproteinase-2
MMP-9: matrix metalloproteinase-9
NAFLD: Non-alcoholic fatty liver disease
NASH: Non-alcoholic steatohepatitis
PPIA: peptidylprolyl Isomerase A
PSR: picrosirius red
ROS: reactive oxygen species
SEAP: secreted alkaline phosphatase
TGF-β1: Transforming growth factor-β 1
TSP-1: Thrombospondin 1
RT-qPCR: real-time quantitative polymerase chain reaction
WT: wild type

