## Supplementary document for "ECM1 Attenuates Hepatic Fibrosis by Interfering with Mediators of Latent TGF-β1 Activation"

### Supplementary Materials and Methods

Table S1. Antibody information

| Antibody | Company | Cat. No. | Dilution |
|---|---|---|---|
| ADAMTS1 | abcam | ab216977 | 1:1000 for WB<br>1:100 for IHC |
| Alexa Fluor 555 goat anti-mouse IgG | Invitrogen | A-21422 | 1:200 |
| Alexa Flour 488 goat anti-rabbit IgG | Invitrogen | A-11008 | 1:200 |
| α-SMA (for IF) | abcam | ab202368 | 1:200 |
| α-SMA (for WB) | abcam | ab7817 | 1:500 |
| α-Tubulin | abcam | ab4074 | 1:1000 |
| Cleaved caspase 3 | Cell signaling technology | 9661 | 1:400 |
| ColI | abcam | ab316222 | 1:500 |
| ColIII | abcam | ab7778 | 1:500 |
| DRAQ5 | Cell signaling technology | 4084L | 1:1000 |
| Anti-human ECM1 | peprotech | 11521-AP | 1:500 for WB;<br>1:100 for IF |
| Anti-mouse ECM1 | abcam | ab253158 | 1:1000 |
| GAPDH | abcam | sc-32233 | 1:1000 |
| HRP-linked anti-mouse | Santa Cruz Biotechnology | sc-2005 | 1:5000 |
| HRP-linked anti-rabbit | Santa Cruz Biotechnology | sc-2357 | 1:5000 |
| MMP-2 (for WB) | Santa Cruz Biotechnology | sc-13595 | 1:500 |
| MMP-2 (for IHC) | Invitrogen | 436000 | 1:100 |
| MMP-9 (for WB) | Santa Cruz Biotechnology | sc-21733 | 1:500 |
| MMP-9 (for IHC) | Invitrogen | MA5-15886 | 1:100 |
| TGF-β1 LAP-D (R58) | Cosmo Bio | RIK-MA-R58 | 1:100 |
| TIMP1 | Santa Cruz Biotechnology | sc-21734 | 1:500 |
| TSP1 | abcam | ab267388 | 1:100 |

Table S2. Primers for RT-qPCR

| Primer | Forward | Reverse |
|---|---|---|
| hADAMTS1 | TTCCACGGCAGTGGTCTAAAG | CCACCAGGCTAACTGAATTACG |
| hACTA2 | GTGTTGCCCCTGAAGAGCAT | GCTGGGACATTGAAAGTCTCA |
| hCOL1A1 | GAGGGCCAAGACGAAGACATC | CAGATCACGTCATCGCACAAC |
| hCOL3A1 | TTGAAGGAGGATGTTCCCATCT | ACAGACACATATTTGGCATGGTT |
| hECM1 | TGAACCAAATCTGCCTTCCTAAC | GCTGGACTGTGGTAGGTTCCA |
| hITGAV | GCTGTCGGAGATTTCAATGGT | TCTGCTCGCCAGTAAAATTGT |
| hITGB6 | GAGGACTACCCGGTGGATTTG | TCCTTTATTGTGTTGAGGTCGTC |
| hMMP-2 | TACAGGATCATTGGCTACACACC | GGTCACATCGCTCCAGACT |
| hMMP-9 | TGTACCGCTATGGTTACACTCG | GGCAGGGACAGTTGCTTCT |
| hPPIA | AGGGTTCCTGCTTTCACAGA | CAGGACCCGTATGCTTTAGG |
| hTGFB1 | AGGGCTACCATGCCAACTTC | CCACGTAGTAGACGATGGGC |
| hTIMP1 | ACCACCTTATACCAGCGTTATGA | GGTGTAGACGAACCGGATGTC |
| hTSP1 | GCCATCCGCACTAACTACATT | TCCGTTGTGATAGCATAGGGG |
| mPpia | GAGCTGTTTGCAGACAAAGTT | CCCTGGCACATGAATCCTGG |
| mEcm1 | GCCAGCTCTGTGGAAGTGGA | CCGGAATCTGTTTATGCTTGC |
| mTsp1 | GGGGAGATAACGGTGTGTTTG | CGGGGATCAGGTTGGCATT |
| mAdamts1 | CATAACAATGCTGCTATGTGCG | TGTCCGGCTGCAACTTCAG |
| mMmp-2 | CAAGTTCCCCGGCGATGTC | TTCTGGTCAAGGTCACCTGTC |
| mMmp-9 | GGACCCGAAGCGGACATTG | CGTCGTCGAAATGGGCATCT |
| mCol1α1 | GCTCCTCTTAGGGGCCACT | CCACGTCTCACCATTGGGG |
| mCol3α1 | CTGTAACATGGAAACTGGGGAAA | CCATAGCTGAACTGAAAACCACC |
| mTimp1 | GCAACTCGGACCTGGTCATAA | CGGCCCGTGATGAGAAACT |
| mActa2 | GTCCCAGACATCAGGGAGTAA | TCGGATACTTCAGCGTCAGGA |

### Human samples

The study protocol was approved by the appropriate local ethics committees (Jing-2015-084 and 2017-584N-MA). Written informed consent was obtained from patients or their representatives. F1-F2 fibrosis liver tissues were collected by liver biopsy. Liver tissue from cirrhotic patients was collected following liver transplantation; allocation and timing of liver transplantation were governed by the China Liver Transplant Registry (CLTR) [1], an official organization for scientific registry authorized by the Chinese Health Ministry, according to individual Model for End-Stage Liver Disease (MELD) scores.

**Mice**

A detailed description of *Ecm1*-knockout (KO) mice can be found in a previous publication [2]. *Fxr*-KO mice were purchased from Jackson Lab. All wild-type (WT), *Ecm1*-KO, and *Fxr*-KO mice were generated from heterozygous predecessors. For KRFK peptide *in vivo* experiments, WT mice received a single tail vein injection of AAV-ECM1 7 days before starting daily intraperitoneal injections of 100 µg KRFK peptide for 14 consecutive days, 100 µg KQFK was used as control. For KTFR peptide *in vivo* experiments, WT and 8-weeks-old Ecm1-KO or *Fxr*-KO mice were injected intraperitoneally with 100 µg KTFR for 14 consecutive days, 100 µg KQFK was used as control. 100 µl of AAV contained 1.25 x $10^{11}$ AAV vector genomes. AAV8-ECM1 was purchased from VectorBuilder. All experiments were conducted with 8- to 10-weeks-old male or female mice. Each group contained 3-6 mice. Animal experiments were performed in compliance with the guidelines for animal care and approved by the local animal welfare committee.

**Cell culture, primary human HSCs isolation, and treatment**

LX-2 HSCs hepatic stellate cells LX-2 HSCs are a human-derived, immortalized HSC cell line first described by Xu et al. [3]. LX-2 HSCs were kept in DMEM supplemented with 1% L-glutamine, 1% P/S and 2% fetal bovine serum (FBS). MFB-F11 is a TGF-β/SMAD reporter cell line that following exposure to TGF-β1,-β2 or-β3 produces secreted alkaline phosphatase (SEAP) and was first described by Tesseur et al. [4]. MFB-F11 cells were grown in DMEM supplemented with 1% P/S and 10% FBS. Primary human HSCs (phHSCs) were isolated by the Cell Isolation Core Facility of the Biobank Großhadern, University Hospital, LMU Munich, using discontinuous density centrifugation with percoll [5]. phHSCs were grown in 10% DMEM supplemented with 1% L-glutamine, 1% P/S and 10% FBS. All cells were kept in a humidified 37°C incubator enriched with 5% $CO_2$.

Following overnight culture, LX-2 HSCs or phHSCs were starved for 4-6h prior to treatment with 100ng/ml recombinant human ECM1 (rECM1), 2 µg/ml recombinant human αvβ6 integrin (3817-AV, R&D Systems, USA), 1 µg/ml recombinant human Thrombospondin-1 (TSP-1) (ECM002, Sigma-Aldrich, USA), 10 ng/ml recombinant human ADAMTS1 (ab134430, abcam, UK), 50 ng/ml recombinant human MMP-2 (ab168864, abcam, UK), 50 ng/ml recombinant human MMP-9 (ab285785, abcam, UK), or 20 µM KTFR peptide for 24-48h, 20 µM KQFK was used as control. For certain experiments, human ECM1 was overexpressed for 24h prior to treatment for 24h in LX-2 HSCs using Lipofectamine® reagents (L3000015, Invitrogen life technologies, USA) according to the manufacturer's instructions followed by treatment with TSP-1, ADAMTS1, MMP-2 or MMP-9 for 24h. MFB-F11 cells were starved for 2h before treatment with conditioned LX-2 HSCs or phHSC supernatant (SN) for 48h or 4 ng/ml latent TGF-β1, 100 ng/ml rECM1 or 20 μM KRFK peptide for 24h, 20 μM KQFK was used as control.

**SEAP activity assay**

Following collection of conditioned media samples from MFB-F11 TGF-β/SMAD reporter cells, SEAP activity and TGF-β concentration were determined using the Great EscAPe® SEAP (631738, Clontech, France) chemiluminescence kit and protocol to quantify SEAP activity (excitation wavelength 360nm; emitted light wavelength 449nm). Chemiluminescence was measured in a Tecan Infinite M200 microplate reader (signal integration time 10000ms) (Tecan Austria, Tecan Group AG, Switzerland). TGF-β concentration was calculated from SEAP activity based on a standard curve of pre-defined TGF-β concentrations (0; 0,1; 0,25; 0,5; 1; 2; 5; 10; 20ng/ml) against SEAP activity which was prepared in advance **(Figure S1A)**.

**Immunoblotting**

Cultured cells were washed 3x with ice-cold phosphate blubbered saline (PBS) and dissolved in RIPA lysis buffer (1% Triton X-100, 50 mMTris [pH 7.5], 300 mM NaCl, 1 mM EGTA, 1 mM EDTA, and 0.1% SDS), supplemented with freshly added phosphatase-protease inhibitors. A DC® Protein Assay (Bio-Rad, USA) was performed according to the manufacturer's protocol to measure sample protein concentration. Quantification was performed in a Tecan Infinite M200® (Tecan Austria, Tecan Group AG, Switzerland) microplate reader using the microplate reader's own protein assay protocol (absorbance at 595nm). Western blot sample protein concentration was calculated using a standard curve of absorbance plotted against pre-defined bovine serum albumin (BSA) concentrations (0, 0.125, 0.25, 0.5, 1, 1.5, 2mg/ml). For each sample, 20 μg of protein were separated by 10% SDS-PAGE gels and were blotted onto a high-resolution nitrocellulose (NC) membrane (MERCK, Darmstadt, Germany). Membranes were blocked with 5% Albumin Bovine Fraction V (SERVA, Heidelberg, Germany) in TBST at room temperature for 1 h. Subsequently, the membrane was incubated with primary antibodies **(Table S1)** overnight at 4°C. The next day, after washing with TBST for 3x, all membranes were incubated with horse radish preroxidase (HRP)-linked anti-mouse or anti-rabbit secondary antibodies. Chemiluminescence was determined in a Fusion® SL chemiluminescence reader (Vilber Lourmat Deutschland GmbH, Germany) from membranes incubated in Western Lightening® Plus-ECL solution (PerkinElmer, USA).

**RNA isolation and real time quantitative (**RT-q)**PCR**

RNA isolation was performed using the TRIzol® reagent (Life Technologies, Carlsbad, CA, USA) according to the manufacturer's instructions. 500 ng RNA were used for cDNA synthesis using a commercially available cDNA synthesis kit (Thermo Scientific, USA). 20 µl mixtures containing 5 µl cDNA (diluted 1:10), 4 µl Power SYBR® Green Master Mix, 10µM forward and reverse primers were used for real time PCR in a StepOnePlus® Real-Time PCR system (Applied Biosystems, Life Technologies, USA). The RT-q PCR amplification protocol comprised a polymerase activation step for 15 minutes at 95 °C, a subsequent amplification step 15s at 95 °C, 20s at 60 °C, and 20s at 72 °C for 40 cycles. A melting curve was established to validate specificity for each PCR analysis with the protocol 15s at 95°C and 1 minute at 60°C, from 60°C to 95°C with +0,3°C every 15 seconds. Human peptidylprolyl isomerase A (hPPIA) was used as a housekeeping gene for normalisation of gene expression. Expressions were calculated with the **ΔΔ**Ct method described previously [6]. Primer sequences were retrieved from the PrimerBank® (Massachusetts General Hospital, USA) online resource and ordered from Eurofins Genomics **(Table S2)**.

**Immunofluorescence (IF) staining and confocal microscopy**

4µm-thick liver slices from formalin-fixed, paraffin-embedded liver tissue were deparaffinized followed by rehydration in xylol and varying concentrations of ethanol (100% and 96%). Antigen unmasking was performed in 1mM EDTA buffer (pH 9.0) using microwave-generated heat for a total of 10 min. Sub-cultured cells were fixed on IF slides with 4% PFA for 10 min. After washing with PBS, liver sections or cell slides were blocked using 0,5% Triton X-100 with 1% BSA in PBS at room temperature for 1h. Subsequently, cells or liver sections were incubated overnight at 4°C with primary anti-TGF-β1 LAP Degradates C-Terminus side cut end R58 (Anti-LAP-R58) (Cosmo Bio, Japan), or anti-ECM1 (11521-1-AP, proteintech, USA) antibody respectively. Subsequently, after washing with PBS, cell slides or liver sections were incubated with Alexa Fluor 555 goat anti-mouse IgG or Alexa Flour 488 goat anti-rabbit IgG (Invitrogen, USA) secondary antibody diluted 1:200 and DRAQ5® Fluorescent Probe Solution diluted 1:1000 in PBS (Thermo Scientific, USA) at room temperature for 1h. After washing with PBS, samples were mounted on IHC/IF glass slides with DakoCytomotion® Fluorescent Mounting Medium (DakoCytomotion, Hamburg, Germany). For confocal microscopy, a Leica DM IRE2 with an HCX PL Apo 63c/1 numeric aperture oil objective (Leica Microsystems, Wetzlar, Germany) was used. The Leica DM IRE2 microscope is a laser scanning spectral confocal microscope. Excitation required a 488nm argon laser, 568nm krypton laser and 633nm HeNe (helium neon) laser. A TCS SP2 scanner was used for image acquisition and Leica Confocal Software for confocal microscopes, version 2.5 (Leica Microsystems, Wetzlar, Germany).

**Immunohistochemistry**

4μm-thick liver slices from formalin-fixed and paraffin-embedded liver tissue were deparaffinized followed by rehydration in xylol and varying concentrations of ethanol (100% and 96%). Antigen unmasking was performed in 1mM EDTA buffer (pH 9.0) using microwave-generated heat for a total of 10 min. Slides were blocked using DAKO Blocking Solution® (S202386-2, Agilent, USA) at room temperature for 30 min. Samples were added one of the following primary antibodies: anti-human LAP-R58 (Cosmo Bio, Japan) , ECM1 (11521-1-AP, proteintech, USA), TSP-1 (18304-1-AP, proteintech, USA), ADAMTS1 (12749-1-AP, proteintech, USA), MMP-2 (436000, Invitrogen, USA), MMP-9 (MA5-15886, Invitrogen, USA) antibodies (diluted 1:100 in PBS); and incubated at 4°C overnight. The next day, all slides were incubated with streptavidin-conjugated horseradish peroxidase antibody at room temperature for 1 hour. Staining was visualized by diaminobenzidine (DAB) and samples were counterstained with hematoxylin as described previously [7]. Slide imaging was performed under a Leica DMRB Leica DMRE® microscope (Leica Microsystems, Wetzlar, Germany) equipped with a Photometrics Coolsnap CCD FX® camera (Leica Microsystems, Wetzlar, Germany). Quantification of target protein staining was performed using the Image J software (National Institute of Health, Bethesda, MD).

**RNAscope assay**

RNAscope was performed according to the manufacturer's instructions using the RNAscope 2.5 HD Detection Kit (Red) (322350, a bio-teche brand, USA). 4 μm-thick liver slices from formalin-fixed and paraffin-embedded (FFPE) liver tissue were deparaffinized followed by rehydration in xylol and varying concentrations of ethanol (100% and 96%). After air-drying for 5 min, all slides were incubated with $H_2O_2$ for 10 min at room temperature. After washing with dd$H_2O$, the slides were boiled in target retrieval buffer (diluted by 10X buffer, 322000, a bio-techne brand, USA) for 30 min. Afterwards, slides were washed twice with dd$H_2O$ and once with 100% ethanol and incubated with Protease Plus for 30 min at 40 °C. After washing again twice with dd$H_2O$ for 2 min, a human *ECM1* probe (RNAscope® Probe- Hs-ECM1, 568211, a bio-teche brand, USA) was added onto the tissue and incubated with the tissue for 2h at 40 °C.

Following incubation, samples were washed twice with $ddH_2O$ for 2 min and placed in 5X sodium saline citrate (SSC) buffer (diluted from 20X stock, 58.44 g/mol sodium chloride, 294.10 g/mol sodium citrate, pH 7) overnight at room temperature. The next day, all slides were incubated with AMP (amplify signal)1-6 reagents for 30 min or 15 min according to the instructions provided by the manual and followed by twice washing with $ddH_2O$ for 2 min respectively. Thereafter, 1:60-diluted Red B and Red A solutions were added for 10 min at room temperature for signal generation. After washing, all samples were counterstained with 50% hematoxylin for 2 min. After washing with $ddH_2O$ for 10 min, slides were dried for 30 min at 37 °C. After dehydration in xylene, slides were mounted using VectaMount (Prolong Gold Antifade Mountant, P10144, Thermo Scientific, USA) and glass coverslips. Slide imaging was performed with Leica DMRB Leica DMRE® microscope (Leica Microsystems, Wetzlar, Germany) equipped with a Photometrics Coolsnap CCD FX® camera (Leica Microsystems, Wetzlar, Germany). Quantification of target protein staining was performed using the Image J software (National Institute of Health, Bethesda, MD).

**Co-immunoprecipitation**

Sub-cultured cells were lysed with RIPA buffer containing 1:100 diluted phosphatase-protease inhibitor cocktail. 1mg of prepared protein was diluted in PBS supplemented with 1:1000 phosphatase-protease inhibitor cocktail to a concentration of 2µg/µl of protein. 100µl of PBS-dissolved 50% protein A/G agarose (sc-2003, Santa Cruz Biotechnology, CA, USA) were added and pre-washed at 4°C for 1h. Following centrifugation at 2000 rpm for 5 min, the resulting supernatant was split into input and Co-IP groups. 125µg IgG or 1µg of anti-ECM1 (11521-1-AP, proteintech, USA) were added to the corresponding samples. Following incubation at 4°C overnight, 100µl of PBS-dissolved 50% protein A/G agarose were added to each sample and incubated again at 4°C overnight. After centrifugation at 2000rpm for 5min, the remaining supernatant was discarded, and the beads were washed with PBS with 1:1000 phosphatase-protease inhibitor cocktail, centrifuged and resuspended in 2x loading buffer of the same volume as protein A/G agarose. Samples were boiled for 5 min prior to SDS-PAGE.

***In vitro* pull-down assay**

1 µg rECM1 and/or 1 µg rTSP-1, rADAMTS1, rMMP-2, or rMMP-9 were incubated with 125 µg IgG or 1 µg ECM1 antibody in 200 µl RIPA buffer containing 1:100 diluted phosphatase-protease inhibitor cocktail at 4°C for 8h. 200 µl of protein A/G agarose were added to the mixture and incubated at 4°C for overnight. Following centrifugation at 2000 rpm for 5 min,

the remaining supernatant was discarded and the beads were washed with PBS (1:1000 phosphatase-protease inhibitor cocktail) three times. After washing and centrifugation, the precipitates were resuspended in 2x loading buffer of the same volume as protein A/G agarose. Samples were boiled for 5 min prior to SDS-PAGE. 10 ng of rECM1 and rTSP-1, rADAMTS1, rMMP-2, or rMMP-9 were loaded as input.

**Matrix metalloproteinase activity assay**

To determine the inhibition of rMMP-2 and -9 (both abcam) activity by rECM1 (R&D Systems), a commercially available fluorescence-based kit was used (ab112146, abcam, UK). Briefly, 10 ng of MMP-2 or -9 were added with 40ng of rECM1 dissolved in assay buffer, incubated for 10 min; followed by the addition of fluorogenic substrate solution and lastly, fluorescence was measured every 5 min for 60 min in a Tecan Infinite M200 microplate reader (excitation wavelength 490 nm; emitted light wavelength 525 nm). rMMP-2 and -9 were purchased as active enzymes and did not require prior activation.

**Computational model**

To estimate the effect of a changed ECM1 concentration during the process of liver injury, repair and fibrosis, the dynamic changes of selected liver cell type fates and molecular signals have been analyzed by simulations with a computational model considering one liver lobule as a well-mixed compartment. Acute liver injury has been studied as a reference. The considered components (cell types, signals) were those considered as highly relevant upon ECM1 concentration changes **(Figure S7)**. Interactions between the components were formulated as "chemical" reactions (R) and translated into a set of ordinary differential equations (ODE) derived from these reactions. The detailed description of reactions and ODE can be found in the supplementary information **(Figure S7, Table S3)**.

**Single cell RNAseq analyses**

The human scRNA-seq dataset (GSE174748) including two normal samples and two NAFLD cirrhosis patient samples was analyzed. Seurat objects were generated from the scRNA-seq gene expression matrix using the Seurat package in R (version 4.4.0). The top 2000 variable genes in each cell scRNA-seq data were identified and normalized using Find Variable Features, Scale Data, and Run PCA functions, which helped to determine the best principal components. Dimensionality reduction was performed using UMAP to effectively summarize these components. The HSCs were isolated from these components and subsequently re-clustered at a resolution of 0.1, resulting in two subclusters. These subclusters of HSCs were then annotated and visualized utilizing the Idents and DimPlot functions, based on established methodologies from previous studies and databases. Heat maps and density plots were utilized to illustrate the expression differences of target genes in HSC subclusters. All clustering and statistical analyses were performed in R (version 4.4.0).

**ELISA**

An ELISA against human TGF-β1 (Invitrogen, USA) was performed according to manufacturer's instructions. 20 μl of LX-2 HSC SN was diluted with 180 μl assay buffer and used for the quantitative detection of human TGF-β1. For the concentration of the total TGF-β1 in the supernatant, 20 μl of 1N HCl was added and mixed to the samples with incubation for 1h at room temperature to activate LTGF-β1 followed by adding 20 μl 1N NaOH for neutralization. For the examination of spontaneous active TGF-β1 in the supernatant, the samples were not treated with HCl and NaOH. Absorbance was measured in a Tecan Infinite M200 microplate reader using 450 nm as the primary wave length (Tecan Austria, Tecan Group AG, Switzerland). TGF-β concentration was calculated based on a standard curve of standardized TGF-β concentrations (0; 0.125; 0.25; 0.5; 1; 2ng/ml) against the absorbance.

**Data availability**

The gene expression omnibus (GEO) DataSets GSE149508 and GSE49541 can be accessed via the following link: https://www.ncbi.nlm.nih.gov/gds. All other data are available in the main text or the supplementary materials. Detailed information not provided within the text will be made available upon request via e-mail to either corresponding author.

**Statistical analysis**

Statistical analyses were performed with GraphPad Prism version 6.0 software. The two-tailed Student's t-test was used to compare two independent groups. One-way ANOVA was adopted to test for statistical differences between the means of two groups. Variables were described by mean and standard deviation (SD). Statistical significance was indicated as follows: *$P < 0.05$; **$P < 0.01$. All experiments were repeated independently at least 3 times.

**Chemical-reaction-inspired schemes:**

***A. Generation of critical molecules:***

$\emptyset\ HH, k_1^{HH} \rightarrow\ ECM1$ (1a)

$\emptyset\ HC, k_1^{HC} \rightarrow\ ECM1$ (1b)

$\emptyset\ qHSC, k_8^{HSC} \rightarrow\ ECM1$ (1c)

$\emptyset\ HH, k_2^{HH} \rightarrow\ LProtease$ (2a)

$\emptyset\ HC, k_2^{HC} \rightarrow\ LProtease$ (2b)

$\emptyset\ HC, k_3^{HC} \rightarrow\ EProtease$ (3a)

$\emptyset\ HH, k_3^{HH} \rightarrow\ EProtease$ (3b)

$\emptyset\ HC, k_4^{HC} \rightarrow\ DAMPs$ (4a)

$\emptyset\ NH, k_1^{NH} \rightarrow\ DAMPs$ (4b)

$\emptyset\ aHSC, k_1^{HSC} \rightarrow\ LProtease$ (5)

$aHSC\ k_0^{HSC} \rightarrow\ aHSC^{(1)}\ k_0^{HSC} \rightarrow\ aHSC^{(2)}\ k_0^{HSC} \rightarrow\ aHSC^{(3)}\ k_0^{HSC} \rightarrow\ aHSC^{(4)}\ k_0^{HSC} \rightarrow$ $\dots aHSC^{(n)}$ (6a)

$\emptyset\ aHSC^{(n)}, k_2^{HSC} \rightarrow\ Matrix,$ (6b)

Where $k_2^{HSC} = k_2^{HSC,0}\left(1 - \left[\frac{V_H + V_S + V_{ECM}}{V_L}\right]\right)$ to take into account that matrix can be produced only as long as there is available physical space i.e., if the lobule volume is not totally absorbed.

$\emptyset\ aHSC, k_3^{HSC} \rightarrow\ LTGF\beta$ (7a)

$\emptyset\ aMph^{+}, k_3^{Mph} \rightarrow\ LTGF\beta,$ (7b)

$k_3^{Mph} = k_{3,0}^{Mph}[aMph^{+}]$, produced by Kupffer cells, assuming, that the production rate of LTGF$\beta$ depends on the presence of Mph Ly6C+ cells.

$\emptyset\ aHSC, k_4^{HSC} \rightarrow\ ProHGF$ (8a)

$\emptyset\ aMph^{+}, k_2^{Mph} \rightarrow\ ProHGF$ (8b)

$\emptyset\ aMph^{+}, k_1^{Mph} \rightarrow\ IFN\gamma$ (9)

$\emptyset\ aMph^{+}, k_4^{Mph} \rightarrow CCL2$ (10a)

$\emptyset\ aIM^{+}, k_4^{Mph} \rightarrow CCL2$ (10b)

$\emptyset\ aHSC, k_7^{HSC} \rightarrow CCL2$ (10c)

All rate constants that have not been explicitly specified are considered constant **(Table S3)**. Eqn (1a), (1b) and (1c) describe the production of $ECM1$ by $HH$, $HC$ and $qHSC$ with the rate $k_1^{HH}$, $k_1^{HC}$ and $k_8^{HSC}$, respectively. Eqn (2a) and (2b) describe the production of $LProtease$ by $HH$ and $HC$ with the rate $k_2^{HH}$ and $k_2^{HC}$, respectively. Eqn (3a) and (3b) describe the production of $EProtease$ by $HC$ and $HH$ with the rate $k_3^{HC}$ and $k_3^{HH}$, respectively. Eqn (4a) and (4b) describe the production of $DAMPs$ by $HC$ and $NH$ with the rate $k_4^{HC}$ and $k_1^{NH}$, respectively. Eqn (5) describes the production of $LProtease$ by $aHSC$ with the rate $k_1^{HSC}$. Eqn (6a) and (6b) describe the process that $aHSC$ first produce some collagen component proteins ($aHSC^{(i)}$ indicates protein-$i$ is produced), which then gradually form the collagen ($Matrix$). Importantly, the introduction of an intermediate reaction, each process being a Poissonian, generates an Erlang distributed waiting time distribution [8], which peaks at a certain value. The more intermediate reactions, the sharper the peak. In general, $V_L$ is the total considered volume of the space (here we consider one liver lobule). $V_H$ is the total volume of all hepatocytes ($HH$, $HC$, $NH$). $V_{ECM}$ is the total volume of the fibrosis-related matrix (denoted by the symbol ECM not to be confused with ECM1). In healthy liver, the volume fraction of hepatocyte, sinusoid, and matrix have been determined to 0.85, 0.11, 0.04 [9]. Hence in healthy steady state, $V_H/V_L$ is 0.85, $V_S/V_L$ is 0.11 and $V_{ECM}/V_L$ is 0.04. Furthermore, we rewrite that $V_H = N_H v_H$, where $N_H$ is the number of hepatocytes, $v_H$ is the average volume of one hepatocyte. We assume the compartment represents one liver lobule x,y-direction with a certain defined height, corresponding to a cylinder of radius of 500 $\mu m$ and the height of 100 $\mu m$. Then $v_H$ is equal to $5.13 \times 10^3\ \mu m^3$ and $V_L$ is equal to $6.49 \times 10^7\ \mu m^3$. The initial number of $N_H$ is about 10760. $V_{ECM} = N_{ECM} v_{ECM}$, where $N_{ECM}$ is the number of moles of collagen, $v_{ECM}$ is one constant parameter equal to $5 \times 10^{18} \mu m^3/mol$ (it is estimated according to the reported molecular mass of collagen, which is 10000 g/mol [10], whereby we assumed the maximal collagen mass density is 2 mg/mL). But since $v_{ECM}$ is too large, which would create numerical instability and influence the robustness of numerical approximation. Therefore, we use another formular $V_{ECM} = M_{ECM} v_{ECM,0}$, where $M_{ECM}$ is the mass of collagen (unit $ng$), $v_{ECM,0}$ is one constant parameter equal to $5 \times 10^5 \mu m^3/ng$. Then the initial value of $M_{ECM}$ is about 5.19 $ng$. In addition, guided by the observation that the sinusoidal network is not influenced by acute drug-induced damage [11], so that $V_S/V_L$ remains 0.11 in the simulations of acute damage, we also approximated the sinusoidal volume by the same value for CLD progression. Eqn (7a) and (7b) describe the production of $LTGF\beta$ by $aHSC$ and $aMph^+$ with rates $k_3^{HSC}$ and $k_3^{Mph}$, respectively. Eqn (8a) and (8b) describe the production of $ProHGF$ by $aHSC$ and $aMph^+$ with rates $k_4^{HSC}$ and $k_2^{Mph}$, respectively. Eqn (9) describes the production of $IFN\gamma$ by $aMph^+$ with rate $k_1^{Mph}$. Eqn (10a) and (10b) describe the production of $CCL2$ by $aMph^+$ and $aHSC$ with rates $k_7^{Mph}$ and $k_4^{HSC}$, respectively.

***B. Reactions between molecular species:***

$$ECM1 + EProtease\ k_1^T \rightarrow\ 0 \qquad (11)$$

$$ECM1 + LProtease\ k_2^T \rightarrow\ [ECM1LProtease] \qquad (12)$$

$$Matrix + LTGF\beta\ k_3^T \rightarrow\ [MatrixLTGF\beta] \qquad (13)$$

$$ECM1 + [MatrixLTGF\beta]\ k_4^T \rightarrow\ [ECM1MatrixLTGF\beta] \qquad (14a)$$

$$ECM1MatrixLTGF\beta\ k_{15}^D \rightarrow MatrixLTGF\beta + ECM1 \qquad (14b)$$

$LProtease + [MatrixLTGF\beta]\ k_5^T \rightarrow\ TGF\beta + Matrix$ (15a)

$LProtease + LTGF\beta\ k_5^T \rightarrow\ TGF\beta$ (15b)

$ProHGF + Matrix\ k_6^T \rightarrow\ [ProHGFMatrix]$ (16)

$ProHGFMatrix + uPA\ k_7^T \rightarrow\ HGF + Matrix$ (17)

All rate constants that have not been explicitly specified are considered constant **(Table S3)**. Eqn (11) describes the consumption of $ECM1$ by $EProtease$. Eqn (12) describes the formation of the complex of $ECM1$ and $LProtease$. Eqn (13) describes the binding of $LTGF\beta$ to $Matrix$. Eqn (14a) and (14b) describe the formation and the dissolution of the complex $ECM1 - MatrixL - TGF\beta$ to mimic the protective role of $ECM1$ on $LTGF\beta$ binding to $Matrix$. Eqn (15a) and (15b) describe the release of $TGF\beta$ from the cleavage of $MatrixLTGF\beta$ and $LTGF\beta$ by $LProtease$. Eqn (16) describes the binding of $ProHGF$ on $Matrix$. Eqn (17) describes the release of $HGF$ from $ProHGF$.

***C. Interactions between cells and molecules:***

$qMph\ k_8^T \rightarrow\ aMph^+,$ (18)

$k_8^T = k_{8,0}^T \frac{[DAMPs]^n}{[DAMPs]^n+[DAMPs_0]^n}$, to take into account that $DAMPs$ is the signal to attract $aMph^+$

$qHSC\ k_9^T \rightarrow\ aHSC$ (19a)

$k_9^T = k_{9,0}^T \frac{[TGF\beta]^n}{[TGF\beta]^n+[TGF\beta_0]^n}$, to take into account that $TGF\beta$ is the signal to activate $qHSC$

$rHSC\ k_{12}^T \rightarrow\ aHSC$ (19b)

$k_{12}^T = k_{12,0}^T \frac{[TGF\beta]^n}{[TGF\beta]^n+[TGF\beta_0]^n}$, to take into account that $TGF\beta$ is the signal to activate $rHSC$

$aHSC\ k_5^{HSC} \rightarrow\ 2aHSC,$ (20)

$k_5^{HSC} = k_{5,0}^{HSC}(1 - \frac{[aHSC]+[qHSC]+[rHSC]}{[HSC_0]})$, to take into account that $aHSC$ can proliferate until the initial number of $HSC$ is reached.

$aHSC\ k_6^{HSC} \rightarrow\ qHSC,$ (21a)

$k_6^{HSC} = k_{6,0}^{HSC}([aMph^-] + [aIM^-]).$

$aHSC\ k_9^{HSC} \rightarrow\ rHSC,$ (21b)

$k_9^{HSC} = k_{9,0}^{HSC}([aMph^-] + [aIM^-]).$

$aHSC\ k_{12}^D \rightarrow\ 0,$ (21c)

$k_{12}^D = k_{12,0}^D([aMph^-] + [aIM^-]).$

$aMph^+\ k_{10}^T \rightarrow\ aMph^-$ (22a)

$k_{10}^T = k_{10}^{T,0}(1 - \frac{[DAMPs]^n}{[DAMPs]^n+[DAMPs_0]^n})$, to take into account that differentiation of $aMph^+$ is enhanced when less $DAMPs$ is present.

$aIM^{+}\ k_{10}^{T} \rightarrow\ aIM^{-}$ (22b)

$aMph^{-}\ k_{11}^{T} \rightarrow\ qMph$ (22c)

$HH\ k_{4}^{HH} \rightarrow\ HC$ (23)

$k_{4}^{HH} = k_{4,0}^{HH}[CCl_4]$.

$HH\ k_{5}^{HH} \rightarrow\ 2HH$ (24)

$k_{5}^{HH} = k_{5}^{HH,0}\left(1 - \left[\frac{V_H+V_S+V_{ECM}}{V_L}\right]\right)$, to take into account that $HH$ can proliferate only as long as there is available physical space i.e., if the lobule volume is not totally absorbed

$HC\ k_{5}^{HC} \rightarrow\ NH$ (25)

$NH\ IM^{+}, k_{5}^{Mph} \rightarrow\ 0$ (26)

$k_{5}^{Mph} = k_{5,0}^{Mph}([aMph^{+}] + [aIM^{+}])$.

$\emptyset\ k_{6}^{Mph} \rightarrow\ aIM^{+}$ (27)

$k_{6}^{Mph} = k_{6,0}^{Mph}[CCL2]$

$aIM^{-}k_{7}^{Mph} \rightarrow\ 0$ (28)

$Matrix + aMph^{-}\ k_{11}^{D} \rightarrow\ aMph^{-}$ (29a)

$Matrix + aIM^{-}\ k_{11}^{D} \rightarrow\ aIM^{-}$ (29b)

All rate constants that have not been explicitly specified are considered constant **(Table S3)**. Eqn (18) describes the activation of $qMph$ by $DAMPs$. Eqn (19a) describes the activation of $qHSC$ by $TGF\beta$. Eqn (19b) describes the activation of $rHSC$ by $TGF\beta$. Eqn (20) describes the proliferation of $aHSC$, where $HSC_0$ is the initial number of HSCs in the compartment. Eqn (21a) describes the switch from $aHSC$ to $qHSC$. Eqn (21b) describes the switch from $aHSC$ to $rHSC$. Eqn (21c) describes the apoptosis of $aHSC$. Eqn (22a) describes the differentiation from inflammatory phenotype of $Mph$ to regenerative phenotype of $Mph$. Eqn (22b) describes the switch from regenerative phenotype of $Mph$ to quiescence. Eqn (23) describes the induced damage to healthy hepatocytes by toxin. Eqn (24) describes the proliferation of healthy hepatocytes. Eqn (25) describes the undergoing necrosis of damaged hepatocytes. Eqn (26) describes the phagocytosis of $NH$ by $aMph^{+}$. Eqn (27) describes the attraction of more $Mph$ by $CCL2$. Eqn (28) describes depletion of $aMph^{-}$. Eqn (29) describes the digestion of $Matrix$ by $aMph^{-}$.

**D. Degradation of molecules**.

$ECM1\ k_{1}^{D} \rightarrow\ 0$

$LProtease\ k_{2}^{D} \rightarrow\ 0$

$EProtease\ k_{3}^{D} \rightarrow\ 0$

$DAMPs\ k_4^D \rightarrow\ 0$

$LTGF\beta\ k_5^D \rightarrow\ 0$

$TGF\beta\ k_6^D \rightarrow\ 0$

$IFN\gamma\ k_7^D \rightarrow\ 0$

$ProHGF\ k_8^D \rightarrow\ 0$

$HGF\ k_9^D \rightarrow\ 0$

$ECM1LProtease\ k_{10}^D \rightarrow\ 0$

$aHSC\ k_{12}^D \rightarrow\ 0,$

$k_{12}^D = k_{12,0}^D[aMph^-]$

$CCl_4\ k_{13}^D \rightarrow\ 0$

$CCL2\ k_{14}^D \rightarrow\ 0$

The degradation processes again maintain mass balance, even if this is not written out explicitly here, as the products of the degradation processes are not tracked. They are approximated for simplicity by first order kinetics. They may in reality follow chains of reactions, higher order reactions or be transported out of the liver compartment, which is not expected to impact on the qualitative dynamics.

**Derived set of ordinary differential equations from the reaction scheme**

The concentration of each molecule and cell at time $t$ in the reaction dynamics can be described as:

$$\partial ECM1(t)/\partial t = k_1^{HC}[HC] + k_1^{HH}[HH] + k_8^{HSC}[HH] + k_{15}^D[ECM1MatrixLTGF\beta] - (k_1^T[Eprotease] + k_2^T[Lprotease] + k_4^T[MatrixLTGF\beta] + k_1^D)[ECM1] \quad (1)$$

where $k_1^{HC} = k_1^{HC,0}(1 - \frac{[IFN\gamma]^n}{[IFN\gamma]^n + [IFN\gamma_0]^n})$.

$$\partial LProtease(t)/\partial t = k_2^{HC}[HC] + k_2^{HH}[HH] + k_1^{HSC}([aHSC] + \sum_i [aHSC^{(i)}]) - (k_2^T[ECM1] + k_5^T[MatrixLTGF\beta] + k_5^T[LTGF\beta] + k_2^D)[LProtease] \quad (2)$$

$$\partial EProtease(t)/\partial t = k_3^{HC}[HC] + k_3^{HH}[HH] - (k_1^T[ECM1] + k_3^D)[EProtease] \quad (3)$$

$$\partial DAMPs(t)/\partial t = k_4^{HC}[HC] + k_1^{NH}[NH] - (k_4^D)[DAMPs] \quad (4)$$

$$\partial LTGF\beta(t)/\partial t = k_3^{HSC}([aHSC] + \sum_i [aHSC^{(i)}]) + k_3^{Mph}[aMph^+] - (k_3^T[Matrix] + k_5^T[LProtease] + k_5^D)[LTGF\beta] \quad (5)$$

$$\partial TGF\beta(t)/\partial t = k_5^T[LProtease][MatrixLTGF\beta] + k_5^T[LProtease][LTGF\beta] - (k_6^D)[TGF\beta] \quad (6)$$

$$\partial INF\gamma(t)/\partial t = k_1^{Mph}[aMph^+] - (k_7^D)[INF\gamma] \quad (7)$$

$$\partial ProHGF(t)/\partial t = k_4^{HSC}([aHSC] + \sum_i [aHSC^{(i)}]) + k_2^{Mph}[aMph^+] - (k_6^T[Matrix] +$$

$k_8^D)[ProHGF]$ (8)

$\partial HGF(t)/\partial t = k_7^T[ProHGFMatrix][uPA] - (k_9^D)[HGF]$ (9)

$\partial ECM1LProtease(t)/\partial t = k_2^T[ECM1][LProtease] - (k_{10}^D)[ECM1LProtease]$ (10)

$\partial MatrixLTGF\beta(t)/\partial t = k_3^T[Matrix][LTGF\beta] + k_{15}^D[ECM1MatrixLTGF\beta] - (k_4^T[ECM1] + k_5^T[LProtease])[MatrixLTGF\beta]$ (11)

$\partial ECM1MatrixLTGF\beta(t)/\partial t = k_4^T[ECM1][MatrixLTGF\beta] - (k_{15}^D)[ECM1MatrixLTGF\beta]$ (12)

$\partial ProHGFMatrix(t)/\partial t = k_6^T[ProHGF][Matrix] - k_7^T[ProHGFMatrix][uPA]$ (13)

$\partial Matrix(t)/\partial t = k_2^{HSC}\left[aHSC^{(n)}\right] + k_5^T[LProtease][MatrixLTGF\beta] + k_7^T[ProHGFMatrix][uPA] - (k_3^T[LTGF\beta] + k_6^T[ProHGF] + k_{11}^D[aMph^-])[Matrix]$ (14), where $k_2^{HSC} = k_2^{HSC,0}\left(1 - \left[\frac{V_H+V_S+V_{ECM}}{V_L}\right]\right)$

$\partial aHSC(t)/\partial t = k_9^T[qHSC] + k_{12}^T[rHSC] + (2k_5^{HSC} - k_6^{HSC} - k_9^{HSC} - k_{12}^D)[aHSC]$ (15), where $k_9^T = k_{9,0}^T\frac{[TGF\beta]^n}{[TGF\beta]^n+[TGF\beta_0]^n}$, $k_{12}^T = k_{12,0}^T\frac{[TGF\beta]^n}{[TGF\beta]^n+[TGF\beta_0]^n}$, $k_5^{HSC} = k_{5,0}^{HSC}\frac{[qHSC]+[rHSC]+[aHSC]}{[HSC_0]}$, $k_6^{HSC} = k_{6,0}^{HSC}([aMph^-] + [aIM^-])$, $k_9^{HSC} = k_{9,0}^{HSC}([aMph^-] + [aIM^-])$, and $k_{12}^D = k_{12,0}^D([aMph^-] + [aIM^-])$

$\partial aHSC^{(i)}(t)/\partial t = k_0^{HSC}\left(\left[aHSC^{(i-1)}\right] - \left[aHSC^{(i+1)}\right]\right) - (k_6^{HSC} + k_{12}^D)[aHSC^{(i)}]$ (16)

$\partial rHSC(t)/\partial t = k_9^{HSC}[aHSC] - k_{12}^T)[rHSC]$ (17)

$\partial qHSC(t)/\partial t = k_6^{HSC}([aHSC] + \sum_i [aHSC^{(i)}]) - k_9^T[qHSC]$ (18)

$\partial aMph^+(t)/\partial t = k_8^T[qMph] - k_{10}^T[aMph^+]$ (19), where $k_8^T = k_{8,0}^T\frac{[DAMPs]^n}{[DAMPs]^n+[DAMPs_0]^n}$ and $k_{10}^T = k_{10}^{T,0}(1 - \frac{[DAMPs]^n}{[DAMPs]^n+[DAMPs_0]^n})$

$\partial aIM^+(t)/\partial t = k_6^{Mph}[CCl_2] - k_{10}^T[aIM^+]$ (20)

$\partial aMph^-(t)/\partial t = k_{10}^T[aMph^+] - k_{11}^T[aMph^-]$ (21)

$\partial aIM^-(t)/\partial t = k_{10}^T[aIM^+] - k_7^{Mph}[aIM^-]$ (22)

$\partial qMph(t)/\partial t = k_{11}^T[aMph^-] - k_8^T[qMph]$ (23)

$\partial CCl_4(t)/\partial t = -k_{13}^D[CCl_4]$ (24)

$\partial CCL2(t)/\partial t = k_4^{Mph}[aMph^+] + k_7^{HSC}([aHSC] + \sum_i [aHSC^{(i)}]) - k_{14}^D[CCL2]$ (25)

$\partial HH(t)/\partial t = 2k_5^{HH}[HH] - k_4^{HH}[CCl_4][HH]$ (26), where $k_5^{HH} = k_5^{HH,0}\left(1 - \left[\frac{V_H+V_S+V_{ECM}}{V_L}\right]\right)$, and $k_4^{HH} = k_{4,0}^{HH}[CCl_4]$

$\partial HC(t)/\partial t = k_4^{HH}[CCl_4][HH] - k_5^{HC}[HC]$ (27)

$$\partial NH(t)/\partial t = k_5^{HC}[HC] - k_5^{Mph}[NH] \quad (28),$$

where $k_5^{Mph} = k_{5,0}^{Mph}([aMph^+] + [aIM^+])$

Table S3. Table of parameter values

| Parameter | Description | Value | Reference |
|---|---|---|---|
| $k_1^{HH}$ | Production rate of $ECM1$ by $HH$ | 0.5e-3 | Estimated |
| $k_2^{HH}$ | Production rate of $LP$ by $HH$ | 1.0e-3 | Estimated |
| $k_3^{HH}$ | Production rate of $EP$ by $HH$ | 1.0e-5 | Estimated |
| $k_4^{HH}$ | Death rate from $HH$ to $HC$ | 5.0e-6 | Estimated |
| $k_5^{HH}$ | Proliferation rate of $HH$ | 1.0e-5 | Estimated |
| $k_1^{HC}$ | Production rate of $ECM1$ by $HC$ | 0.5e-4 | Estimated |
| $k_2^{HC}$ | Production rate of $LP$ by $HC$ | 1.0e-2 | Estimated |
| $k_3^{HC}$ | Production rate of $EP$ by $HC$ | 1.0e-4 | Estimated |
| $k_4^{HC}$ | Production rate of $DAMPs$ by $HC$ | 1.0e-4 | Estimated |
| $k_5^{HC}$ | Death rate from $HC$ to $NH$ | 1.0e-5 | Estimated |
| $k_1^{NH}$ | Production rate of $DAMPs$ by $NH$ | 1.0e-3 | Estimated |
| $k_0^{HSC}$ | Switch rate from $aHSC^{(i)}$ to $aHSC^{(i+1)}$ | 1.0e-4 | Estimated |
| $k_1^{HSC}$ | Production rate of $LP$ by $aHSC$ | 1.0e-6 | Estimated |
| $k_2^{HSC}$ | Production rate of $fMatrix$ by $aHSC$ | 4.0e-5 | Estimated |
| $k_3^{HSC}$ | Production rate of $LTGF\beta$ by $aHSC$ | 1.0e-4 | Estimated |
| $k_4^{HSC}$ | Production rate of $proHGF$ by $aHSC$ | 1.0e-4 | Estimated |
| $k_5^{HSC}$ | Proliferation rate of $aHSC$ | 3.4e-5 | Estimated |
| $k_6^{HSC}$ | Switch rate from $aHSC$ to $qHSC$ | 1.0e-5 | Estimated |
| $k_7^{HSC}$ | Production rate of $CCL2$ by $aHSC$ | 1.0e-4 | Estimated |
| $k_8^{HSC}$ | Production rate of $ECM1$ by $qHSC$ | 1.0e-6 | Estimated |
| $k_9^{HSC}$ | Switch rate from $aHSC$ to $rHSC$ | 1.0e-5 | Estimated |
| $k_1^{Mph}$ | Production rate of $IFN\gamma$ by a$Mph+$ | 1.0e-4 | Estimated |
| $k_2^{Mph}$ | Production rate of $proHGF$ by a$Mph+$ | 1.0e-4 | Estimated |
| $k_3^{Mph}$ | Production rate of $LTGF\beta$ by $aMph+$ | 1.0e-3 | Estimated |
| $k_4^{Mph}$ | Production rate of $CCL2$ by $aMph+$ | 1.0e-3 | Estimated |
| $k_5^{Mph}$ | Degradation rate of $NH$ | 1.0e-5 | Estimated |
| $k_6^{Mph}$ | Attraction rate of $aMph+$ by $CCL2$ | 12.0e-6 | Estimated |

| | | | |
|---|---|---|---|
| $k_7^{Mph}$ | Degradation rate of $aMph-$ | 1.0e-4 | Estimated |
| $k_1^T$ | Degradation rate of $ECM1$ by $EP$ | 1.0e-4 | Estimated |
| $k_2^T$ | Degradation rate of $ECM1$ by $LP$ | 1.0e-1 | Estimated |
| $k_3^T$ | Binding rate of $LTGF\beta$ to $Matrix$ | 1.0e-3 | Estimated |
| $k_4^T$ | Binding rate of $ECM1$ to $MatrixLTGF\beta$ | 1.0e-6 | Estimated |
| $k_5^T$ | Binding rate of $LP$ to $MatrixLTGF\beta$ | 1.0e-7 | Estimated |
| $k_6^T$ | Binding rate of $proHGF$ to $Matrix$ | 1.0e-3 | Estimated |
| $k_7^T$ | Releasing rate of $HGF$ from $proHGFMatrix$ | 1.0e-3 | Estimated |
| $k_8^T$ | Activation rate from $qMph$ to $aMph+$ | 1.0e-3 | Estimated |
| $k_9^T$ | Activation rate from $qHSC$ to $aHSC$ | 1.0e-4 | Estimated |
| $k_{10}^T$ | Differentiation rate from $aMph+$ to $aMph-$ | 1.0e-5 | Estimated |
| $k_{11}^T$ | Switch rate from $aMph-$ to $qMph$ | 1.0e-3 | Estimated |
| $k_{12}^T$ | Activation rate from $rHSC$ to $aHSC$ | 1.0e-2 | Estimated |
| $k_1^D$ | Degradation rate of $ECM1$ | 1.0e-5 | Xiao & Wu 2017 [12] |
| $k_2^D$ | Degradation rate of $EP$ | 2.4e-5 | McGray et al. 2011 [13]; O'Sullivan et al. 2014 [14]; Takeuchi et al. 2023 [15] |
| $k_3^D$ | Degradation rate of $LP$ | 2.4e-5 | McGray et al. 2011 [13]; O'Sullivan et al. 2014 [14]; Takeuchi et al. 2023 [15] |
| $k_4^D$ | Degradation rate of $DAMPs$ | 5.7e-4 | Zandarashvili et al. 2013 [16] |
| $k_5^D$ | Degradation rate of $LTGF\beta$ | 1.2e-4 | Wakefield et al. 1990 [17] |
| $k_6^D$ | Degradation rate of $TGF\beta$ | 5.8e-3 | Wakefield et al. 1990 [17] |
| $k_7^D$ | Degradation rate of $IFN\gamma$ | 3.8e-4 | Miyakawa et al. 2011 [18] |
| $k_8^D$ | Degradation rate of $proHGF$ | 5.8e-3 | Chang et al. 2016 [19] |
| $k_9^D$ | Degradation rate of $HGF$ | 5.8e-3 | Chang et al. 2016 [19] |
| $k_{10}^D$ | Degradation rate of $ECM1LP$ | 2.4e-5 | McGray et al. 2011 [13]; O'Sullivan et al. 2014 [14]; Takeuchi et al. 2023 [15] |
| $k_{11}^D$ | Degradation rate of $fMatrix$ by $aMph-$ | 1.2e-4 | Estimated |
| $k_{12}^D$ | Degradation rate of $aHSC$ by $aMph-$ | 1.0e-5 | Estimated |

| | | | |
|---|---|---|---|
| $k_{13}^{D}$ | Degradation rate of $CCL4$ | 6.4e-5 | Estimated |
| $k_{14}^{D}$ | Degradation rate of $CCL2$ | 5.8e-4 | Berchiche et al. 2011 [20] |
| $k_{15}^{D}$ | Decomposition rate of $ECM1MatrixLTGF\beta$ into $MatrixLTGF\beta$ and $ECM1$ | 1.0e-5 | Estimated |

The degradation rate of each signal (e.g. $k_1^D$ for $ECM1$) is calculated according to its half-life time: $2\ /T_{1/2,ECM1}$. The production rate of each signal is not based on real biological data, which is difficult to measure by experiment. Therefore, the value simply represents a qualitative estimation of the production. The death rate ($k_4^{HH}$) and proliferation rate ($k_5^{HH}$) of hepatocyte are estimated to fit the observation that about half of the hepatocytes are killed by acute $CCl_4$ injection and then restore after about 6 days [11]. The differentiation rate ($k_0^{HSC}$) of aHSC is to fit the observation that α-SMA positive HSC is detected one day after $CCl_4$ injection [21]. The degradation rates of aHSC ($k_{12}^{D}$), aMph ($k_7^{Mph}$), and fibrotic matrix ($k_{11}^{D}$) are estimated to make sure they restore to the initial value after about one week.

**Supplementary figures and figure legends**

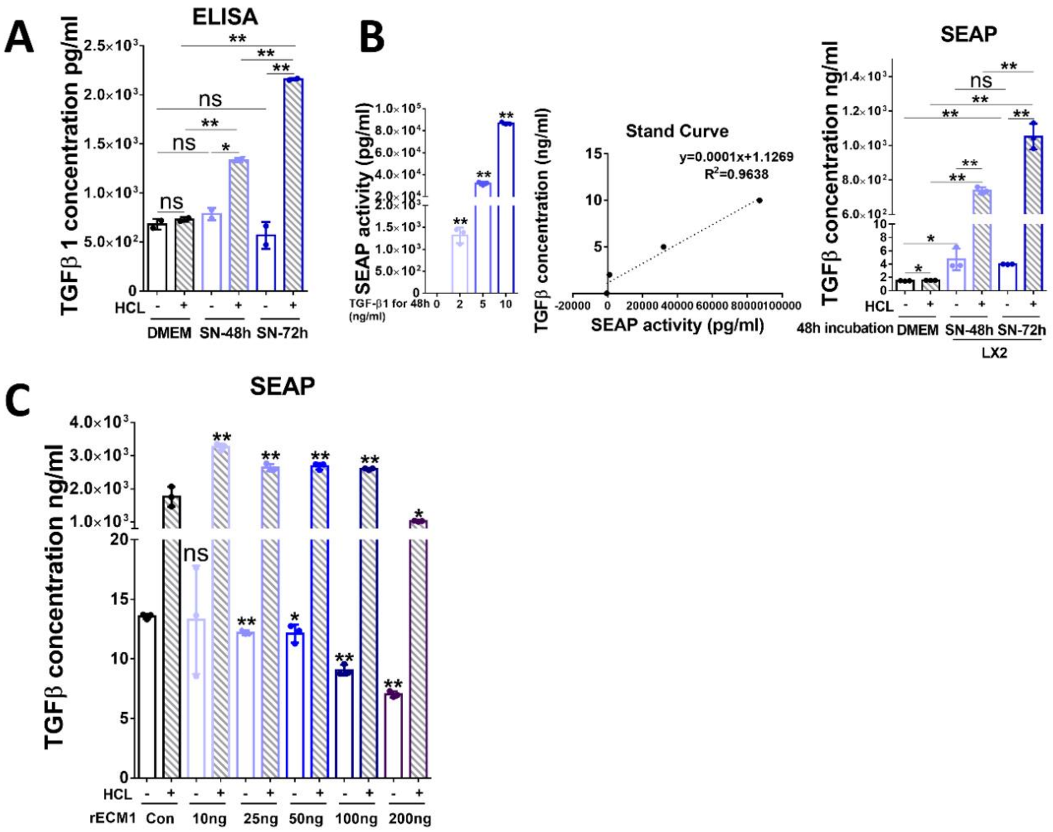


**Figure S1: Establishment of SEAP activity assay**

(A) The concentration of active and total TGF-β1 in DMEM (control) and LX-2 SN collected after 48h and 72h of incubation were examined by human TGF-β1 ELISA.

(B) TGF-β1 standard curve was calculated by SEAP-activity of MFB-F11 cells in response to the treatment with different concentrations of TGF-β1 (0, 2, 5, 10 ng/ml) for 48h (left and middle); Active and total (with HCl treatment) TGF-β concentration (ng/ml) in DMEM (control) and LX-2 SN collected after 48h and 72h of incubation were tested by SEAP activity assay with the standard curve shown above.

(C) Active and total (with HCl treatment) TGF-β concentration (ng/ml) were examined by SEAP activity assay in LX-2 SN following treatment with increasing concentrations of rECM1 for 48h.

P values were calculated by unpaired Student's t-test. Bars represent mean±SD. *p<0.05; **p<0.01.

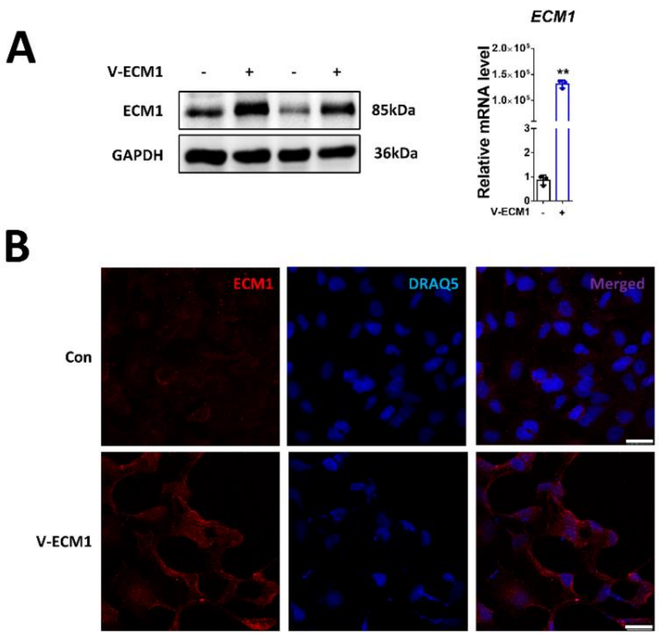


**Figure S2: Human ECM1 plasmid overexpression in LX-2 HSCs**

(A) RT-qPCR and Western blot of human ECM1 in LX-2 HSCs following treatment with human ECM1 plasmid (V-ECM1). For RT-qPCR, *PPIA* was used as endogenous control. P values were calculated by unpaired Student's t-test. Bars represent mean ± SD. *p<0.05; **p<0.01. For Western blotting, GAPDH was used as a loading control.

(B) Immunofluorescence of human ECM1 in LX-2 HSCs following treatment with human ECM1 plasmid (V-ECM1). DRAQ5 was used for nuclear staining. Scale bar, 25 µm.

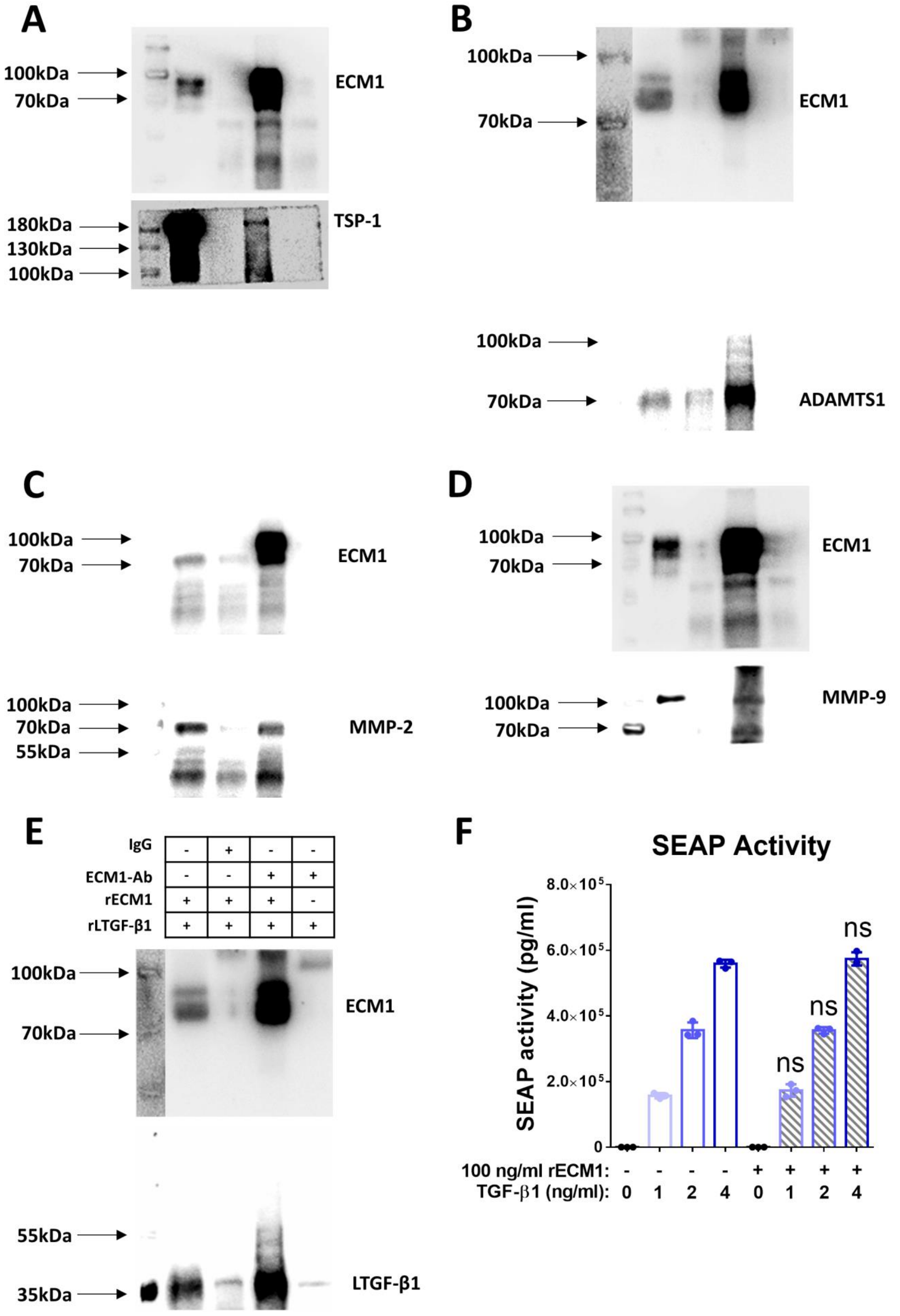


**Figure S3. In vitro binding assay of rECM1 and rTSP-1, rADAMTS1, rMMP-2, rMMP-9, rLTGF-β1, and active TGF-β1.**

(A-E) In vitro pull-down assays of rECM1 and rTSP-1, rADAMTS1, rMMP-2, rMMP-9, rLTGF-β1.

(F) The SEAP activity (ng/ml) from conditioned MFB-F11 supernatant incubated with 100ng/ml rECM1 and/or 0, 1, 2, 4 ng/ml active TGF-β1. P values were calculated by unpaired Student’s t-test. Bars represent mean ± SD. *p<0.05; **p<0.01.

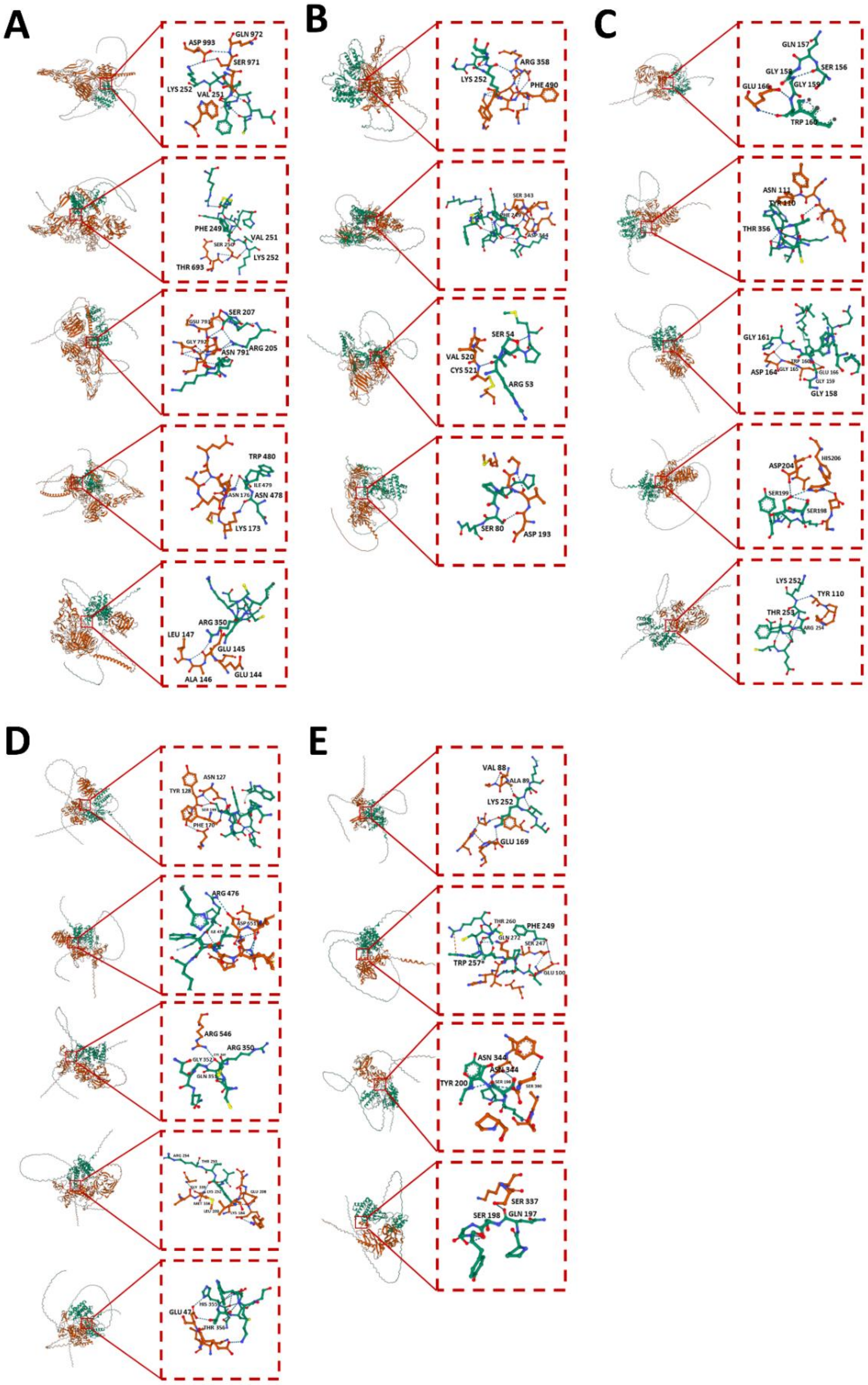

**Figure S4. AlphaFold structural modeling of the ECM1 and TSP-1 (A), ADAMTS1 (B), MMP-2 (C), MMP-9 (D), or LTGF-β1 (E) interaction interface.**

4 or 5 models of the each interaction interface generated by AlphaFold3, with full-length of the target proteins as the input sequences. The ECM1 protein and residues involved in the interaction were shown in green in five figures while TSP-1, ADAMTS1, MMP-2, MMP-9, or LTGF-β1 were shown in red from A to E respectively. Opened with Visual Studio Code and edited with Protein Viewer V0.1.0.

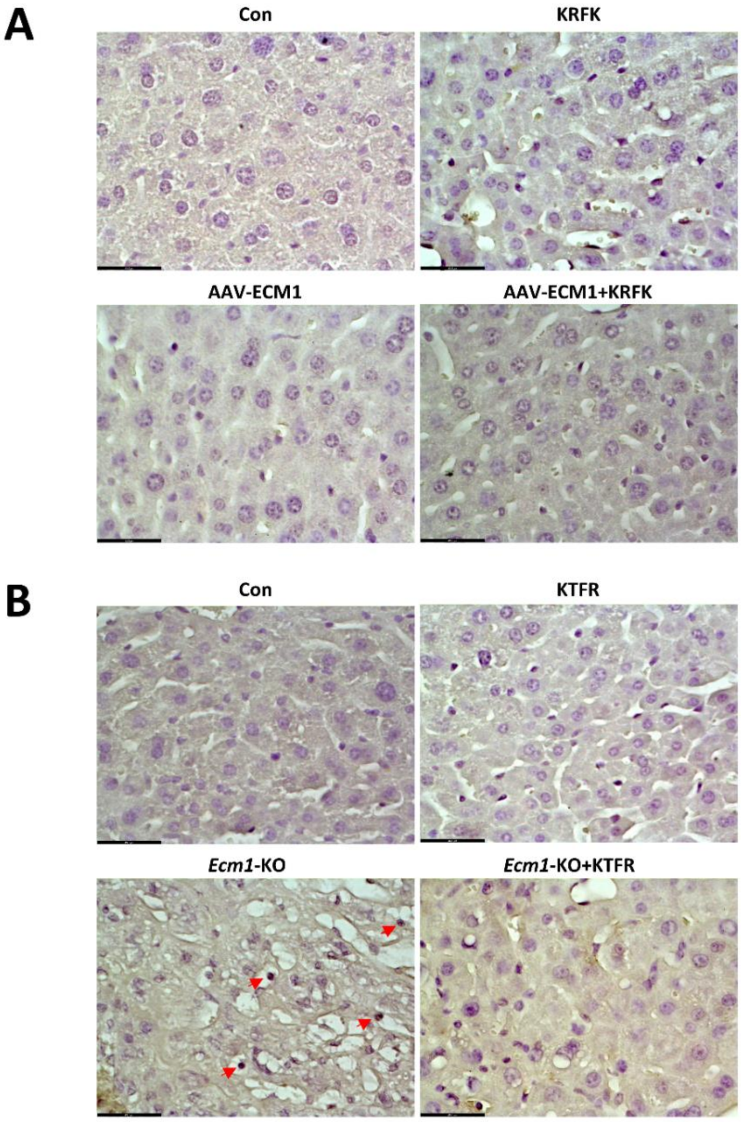


**Figure S5. Cell apoptosis analyses in liver tissue from WT or *Ecm1*-KO mice with or without AAV-ECM1 or KQFK/KRFK/KTFR treatment.**

(A) Representative images of IHC staining for cleaved caspase 3 in liver tissue from WT mice treated with or without AAV-ECM1 or KQFK/KRFK. Scale bar, 43.5 μm.

(B) Representative images of IHC staining for cleaved caspase 3 in liver tissue from WT or *Ecm1*-KO mice treated with KQFK or KTFR peptides. Arrows indicate the positive nuclear staining of cleaved caspase 3. Scale bar, 43.5 μm.

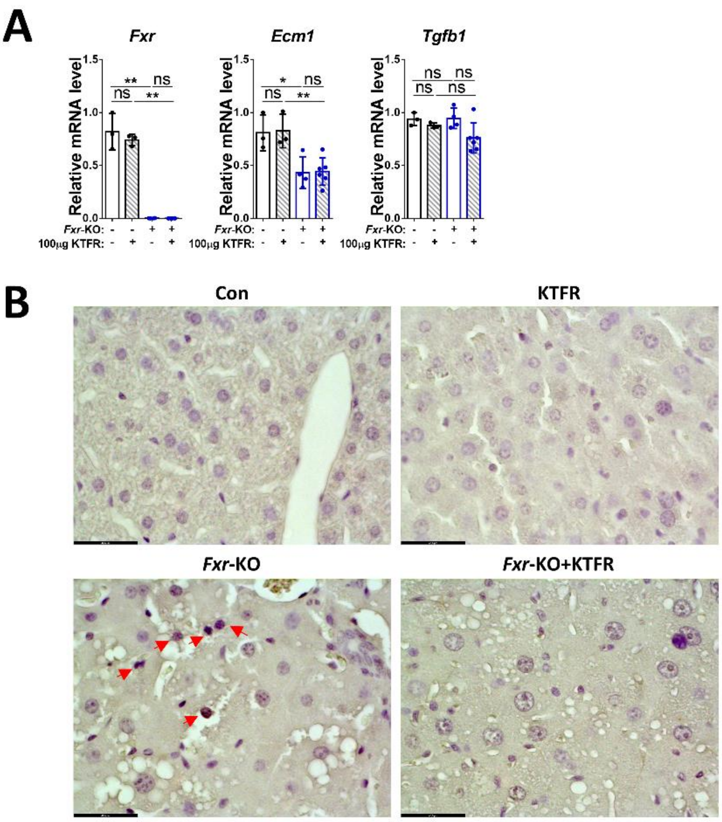


**Figure S6. Relative mRNA expression and cell apoptosis analysis in liver tissue from WT or *Fxr*-KO mice injected with KQFK or KTFR peptides.**

(A) RT-qPCR of *Fxr*, *Ecm1*, and *Tgfb1* in liver tissue from WT or *Fxr*-KO mice injected with KQFK or KTFR peptides. *PPIA* was used as endogenous control. P values were calculated by unpaired Student's t-test. Bars represent mean ± SD. *p<0.05; **p<0.01.

(B) Representative images of IHC staining for cleaved caspase 3 in liver tissue from WT or *Fxr*-KO mice treated with KQFK or KTFR peptides. Arrows indicate the positive nuclear staining of cleaved caspase 3. Scale bar, 43.5 μm.

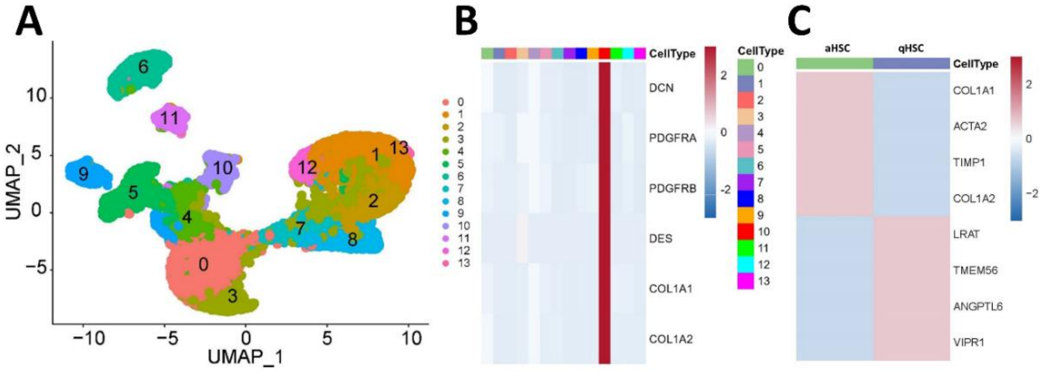


**Figure S7. The scRNA-seq analysis of HSCs in controls and NAFLD cirrhotic patients.**

(A) UMAP visualization of cell clusters from controls and NAFLD cirrhotic patients.

(B, C) Heatmap illustrating the marker genes utilized for annotating HSCs (B) and HSCs subclusters.

### Computational model of ECM1 pathway network

**Model structure**. The dynamic changes of selected liver cell type fates and signals during the process of injury, repair and fibrosis is analyzed by a compartment model that executes a series of reactions (R) derived from the sketch, as shown in **Figure S7**. These cells and signals are considered: quiescent HSCs (qHSC), reverted HSCs (rHSC), activated HSCs (aHSC), resident macrophages of appropriate phenotype denoted as aMph (Ly6C-high phenotype as $aMph^{+}$, Ly6C-low phenotype as $aMph^{-}$), infiltrating macrophages of appropriate phenotype denoted as aIM (Ly6C-high phenotype as $aIM^{+}$, Ly6C-low phenotype as $aIM^{-}$), healthy hepatocytes (HH), damaged hepatocytes (HC), necrotic hepatocytes (NH), ECM1, L-Mediator, E-Protease, LTGF-β1, TGF-β1, ProHGF, HGF, DAMPs, CCL2, IFNγ. The compartment model considers a liver lobule as a well-mixed (stirred) reaction and does not resolve the positions of species (cells or signal molecules) in space.

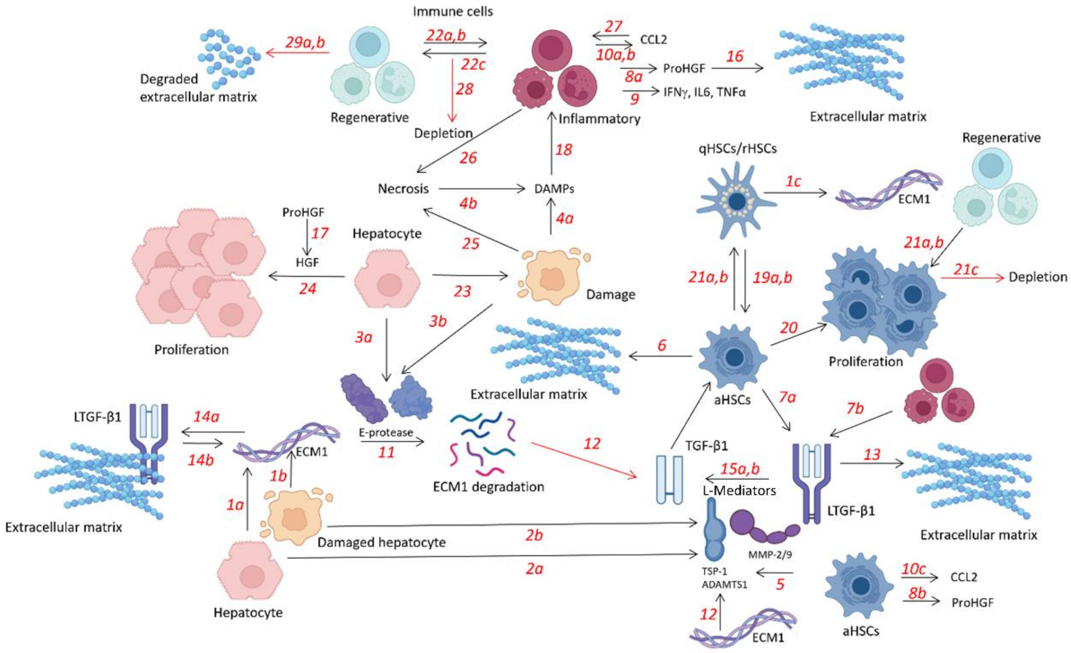


**Figure S8**. **Scheme comprising all interactions in the compartment model of liver injury.** It includes the major molecules produced by corresponding cell types upon liver injury and their interactions to regulate the recovery process that follows. Detailed description for each arrow, representing one interaction or process, can be found under model structure: Arrows 1—10 are described in section A representing the generation of critical molecules by cells; arrows 11—17 are described in section B, displaying interactions between molecular species; arrows 18—29 are described in section C and specify interactions between cells and molecules.

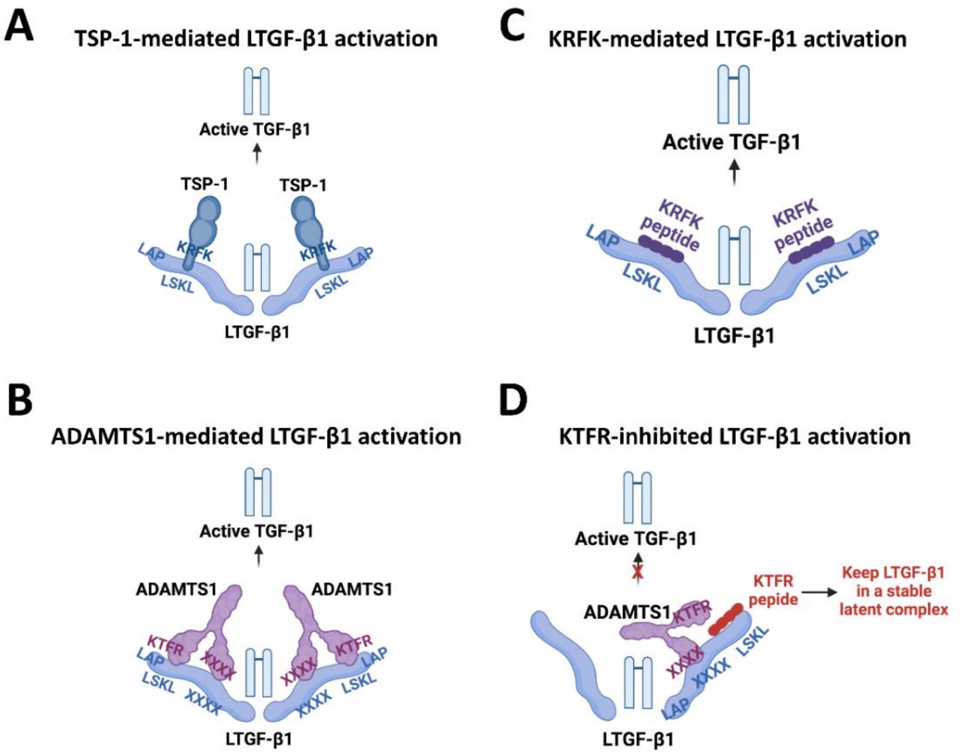


**Figure S9. Scheme showing LTGF-β1 activation mediated by (A) TSP-1, (B) ADAMTS1, (C) KRFK peptide, and (D) KTFR peptide.**

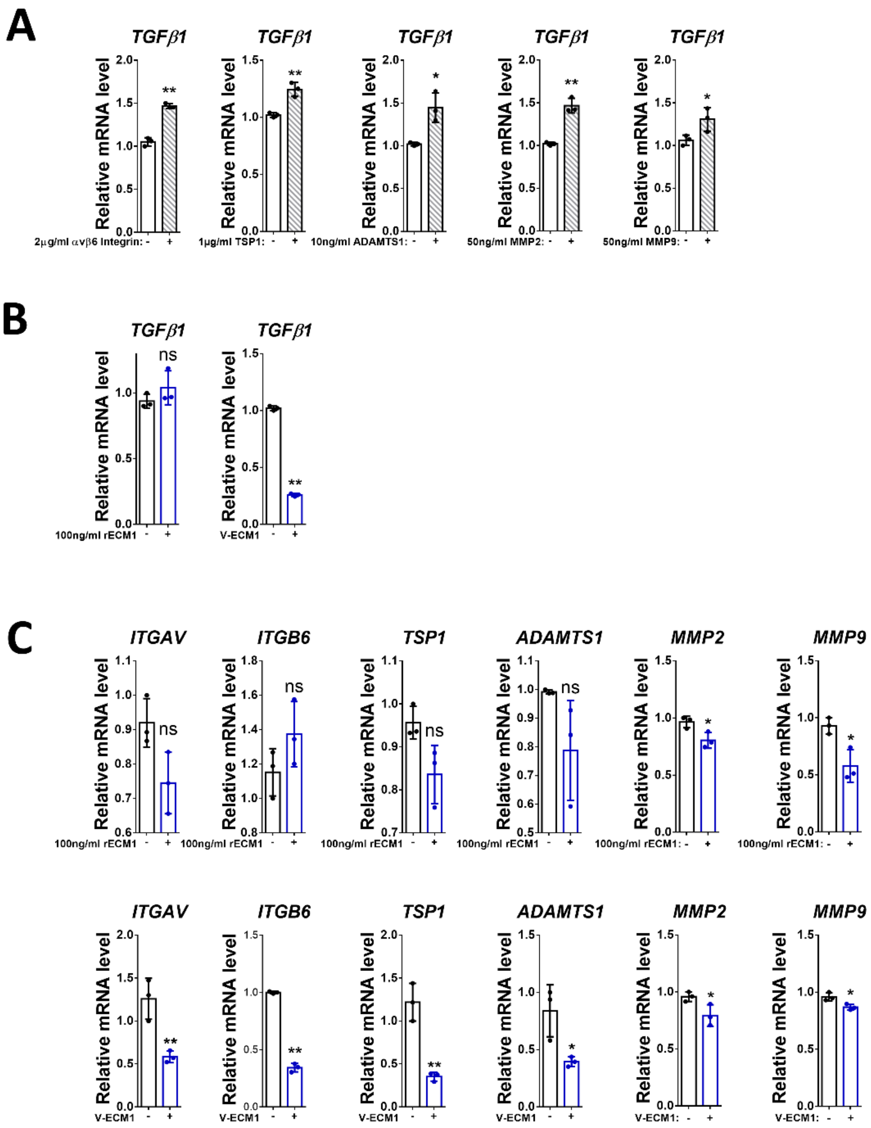


**Figure S10. Effects of LTGF-β1 activators and ECM1 on mRNA expression TGF-β1; effects of ECM1 on mRNA expression of LTGF-β1 activators in LX-2 HSCs.**

(A) RT-qPCR of *TGFβ1* mRNA expression in LX2 HSCs treated with αvβ6 Integrin, TSP-1, ADAMTS1, MMP-2, or MMP-9.

(B) RT-qPCR of *TGFβ1* mRNA expression in LX2 HSCs incubated with ECM1 or transfected with V-ECM1.

(C) RT-qPCR of *ITGAV*, *ITGB6*, *TSP-1*, *ADAMTS1*, *MMP2*, and *MMP9* in LX-2 HSCs incubated with ECM1 or transfected with V-ECM1.

For RT-qPCR, *PPIA* was used as endogenous control. P values were calculated by unpaired Student's t-test. Bars represent mean ± SD. *p<0.05; **p<0.01.

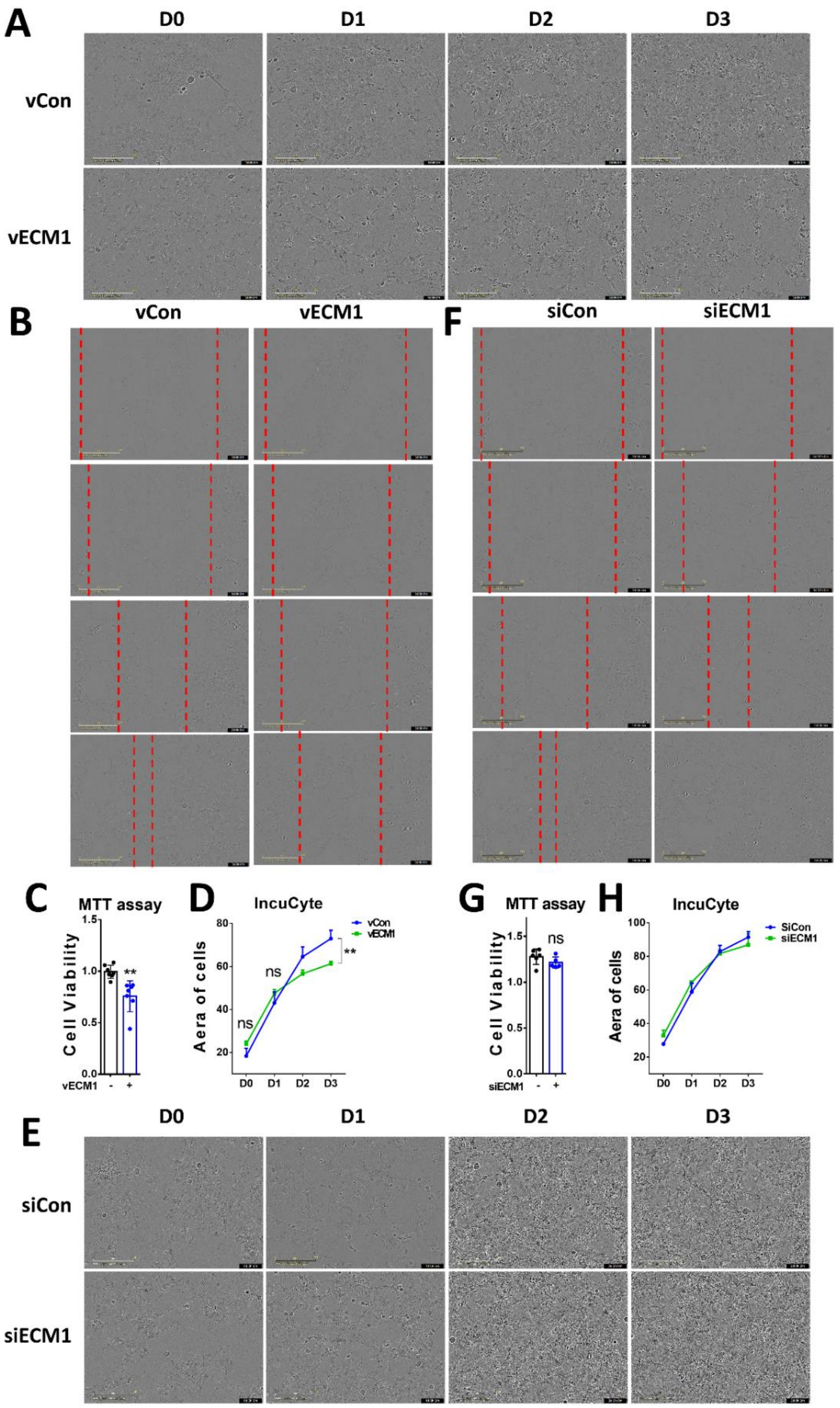


**Figure S11. Effects of ECM1 on HSC proliferation and migration.**

(A)(E) Bright field photos of LX-2 cells treated with control, ECM1 overexpression or knockdown.

(B) (F) Bright field photos of LX-2 cells showing migration assay mediated by ECM1 overexpression or knockdown.

(C) (G) MTT analyses showing LX-2 cell proliferation with control, ECM1 overexpression, or knockdown treatment.

(D) (H) IncuCyte assay of LX-2 cell proliferation with control, ECM1 overexpression, or knockdown treatment.

18 Miyakawa N, Nishikawa M, Takahashi Y, Ando M, Misaka M, Watanabe Y*, et al.* Prolonged circulation half-life of interferon gamma activity by gene delivery of interferon gamma-serum albumin fusion protein in mice.

J Pharm Sci 2011;**100**:2350-7.
19 Chang HK, Kim PH, Cho HM, Yum SY, Choi YJ, Son Y, *et al.* Inducible HGF-secreting Human Umbilical Cord Blood-derived MSCs Produced via TALEN-mediated Genome Editing Promoted Angiogenesis. Mol Ther 2016;**24**:1644-54.
20 Berchiche YA, Gravel S, Pelletier ME, St-Onge G, Heveker N. Different effects of the different natural CC chemokine receptor 2b ligands on beta-arrestin recruitment, Galphai signaling, and receptor internalization. Mol Pharmacol 2011;**79**:488-98.
21 Jieling Zhao AG, Reham Hassan, Steven Dooley, Jan Georg Hengstler, Dirk Drasdo A liver digital twin for in silico testing of cellular and inter-cellular mechanisms in regeneration after drug-induced damage. iScience 2023.
